# De novo rates of a *Trypanosoma*-resistant mutation in two human populations

**DOI:** 10.1101/2024.10.10.617206

**Authors:** Daniel Melamed, Revital Shemer, Evgeni Bolotin, Michael B. Yakass, Dorit Fink-Barkai, Edem K. Hiadzi, Karl L. Skorecki, Adi Livnat

## Abstract

Until recently, mutation rates have only been measured as averages across many genomic positions. Recently, a method to measure mutation rates at the single-mutation resolution was applied to a narrow region in the human hemoglobin subunit beta (*HBB*) gene containing the site of the hemoglobin S (HbS) mutation as well as to a paralogous hemoglobin subunit delta (*HBD*) region, in sperm samples from sub-Saharan African and northern European donors. The HbS mutation, which protects against malaria while causing sickle-cell anemia in homozygotes originated de novo significantly more frequently in the *HBB* gene in Africans compared to the other three test cases combined (the European *HBB* gene and the European and African *HBD* gene). Here, we apply this approach to the human apolipopro-tein L1 (*APOL1*) gene containing the site of the G1 1024A*→*G mutation, which protects against African sleeping sickness caused by *Trypanosoma brucei gambiense* while causing a substantially increased risk of chronic kidney disease (CKD) in homozygotes. We find that the 1024A*→*G mutation is the mutation of highest de novo origination rate and deviates most from the genome-wide average rate for its type (A*→*G) compared to all other observable mutations in the region, and that it originates de novo significantly more frequently in Africans than in Europeans—i.e., in the population where it is of adaptive significance. The results are inconsistent with the notion that the probability of a specific mutational event is independent of its value to the organism and underscore the importance of studying mutation rates at the single-mutation resolution.

## Introduction

Until recently, technical limitations have prevented researchers interested in studying mutation rates from examining anything other than mutation rates averaged across many genomic positions. For example, researchers have estimated the average mutation rate across the *∼*3 billion base pairs of the entire haploid human genome (*1–3*), the average mutation rate across the many instances of any given motif (*3–5*) (e.g., the A*→*G mutation rate in the middle of the AAG motif, estimated from the frequency of extremely rare variants averaged across nearly 29,000 instances of this motif per diploid human genome and across nearly 4,000 individuals (*5*)), or the mutation rate across the stretch of a gene in certain cases (*6–8*) (e.g., averaged across the 3,657 bp of the locus implicated in Alagille Syndrome, AGS (*8*)). Such averages combine observations on many positions in the genome into one summary statistic, which, while informative, precludes a far more precise assessment of mutation rates of individual mutations at individual base positions. For instance, while it is known that the C*→*T mutation occurs more frequently in the CpG motif than it occurs on average across the whole genome (*9–11*), this does not tell us what the rate of the C*→*T mutation is in any one particular instance of the motif in any particular position in the genome and whether it is higher or lower than the rate of the same mutation in any other motif-instance in any other position.

Recently, Melamed et al. (*12*) developed a method that combines controlled mutation enrichment by a restriction enzyme and a deep sequencing–based approach that for the first time had enough accuracy and yield to measure the de novo rates of individual mutations at individual base positions. They applied this method to a 6 base pair region of interest (ROI) in the human hemoglobin subunit beta gene (*HBB*) and to the identical paralogous region of the hemoglobin subunit delta gene (*HBD*) (*12*). The *HBB* region includes the site of the 20A*→*T mutation— commonly referred to as the hemoglobin S (HbS) mutation—which protects against malaria while causing sickle cell anemia in homozygotes (*13–16*), whereas *HBD* mutations have not been impli-cated in malaria resistance (*17*). These genomic regions were examined in human sperm samples from both sub-Saharan and northern European donors (*12*). Results showed, first, that the rates of the same mutations varied substantially between the two genes and the two populations even though these mutations appeared on the same local genetic background in these genes and populations (*12*), and that the population-level difference in the point mutation rate averaged across the 6 bp *HBB* region was two orders of magnitude larger than previously measured population-level differences in mutation rates averaged across many loci (*12, 18, 19*). Given that this was the first examination of the rates of individual mutations and that the mutation rate variation obtained was notably higher than that of mutation rate averages across many positions obtained in previous studies, could it be that an important part of the adaptively meaningful variation in mutation rates is to be found at this resolution of individual mutations?

In addition to demonstrating high mutation rate variation at the individual-mutation resolution for the first time, results showed that the 20A*→*T mutation originated de novo significantly more frequently in the *HBB* gene in sub-Saharan African donors, where it is adaptively significant, compared to the three cases where it is not, combined: the *HBB* gene in European donors and the *HBD* gene in both African and European donors (*12*). That is, given that malaria is common in sub-Saharan Africa and that the 20A*→*T mutation is protective against it when it occurs in *HBB* but not in *HBD*, this mutation was found to originate de novo significantly more frequently in the gene and in the population where it is of adaptive significance (*12*). This finding was unexpected given the central tenet of neo-Darwinism that mutations are random, where this randomness “refers to the supposition that the likelihood of any particular mutational event is independent of its specific value to the organism” (*20*). Given that the probabilities of particular mutational events had not been measured before, could it be that there actually is a relationship between the likelihood of a particular mutational event and its specific value to the organism, which could not have been detected with previous methods?

It is tempting to answer with a resounding “no” to this question, given that there could hardly be a more fundamental assumption in evolutionary theory that data could violate (*21*). Yet the only way of actually knowing the answer is to carry out more studies of the probabilities of particular mutational events. Here, we examined a different gene and mutation that has also been directly implicated in adaptation: the human apolipoprotein L1 (*APOL1*) G1 1024A*→*G mutation.

APOL1 is the only known member of the apolipoprotein L family of innate immunity factors which is secreted into the bloodstream, where it associates with serum High Density Lipoprotein-C particles. These, in turn, carry the “good cholesterol” and exhibit anti-inflammatory activity (*22*). Human APOL1 has also evolved to confer protection against an extended spectrum of *Trypanosoma brucei* species, spread by the *Glossina* species of the Tse Tse fly vector, which cause African sleeping sickness when unchecked—a devastating disease of wide historical prevalence and impact in Africa (*23*). In what appears to be an evolutionary arms race, two *Trypanosoma brucei* species—*Trypanosoma brucei gambiense* in West Africa and *Trypanosoma brucei rhodesiense* in East Africa (*24–28*)—evolved mechanisms of resistance to the ancestral human APOL1, and two sets of human APOL1 variants evolved in Africa, each providing protection respectively against one of these species (*29, 30*). The G1 variants comprise two non-synonymous mutations in nearly perfect linkage disequilibrium: rs73885319 (1024A*→*G), altering codon 342 from Serine to Glycine (S342G), and rs60910145 (1152T*→*G), altering codon 384 from Isoleucine to Methionine (I384M), and it has been shown that both homozygotes and heterozygotes for the first of these mutations (1024A*→*G) are asymptomatic for *T. b. gambiense* (*26*). The G2 variant (rs71785313)—consisting of a 6 nucleotide in-frame deletion of two adjacent codons for Aspargine and Tyrosine at positions 388 and 389—protects against *T. b. rhodesiense:* neither homozygotes nor heterozygotes become infected (*26*). Besides these protective abilities, however, the G1/G1 and G2/G2 homozygotes as well as the G1/G2 compound heterozygote impose a substantially increased risk of chronic kidney disease (CKD) (*29, 30*)—itself a devastating condition affecting millions of people of African descent worldwide (*31*), though there has been rapidly emerging progress in developing precision therapeutics (*32*). The G1 and G2 variants are prevalent in populations that live in or trace their recent ancestry to sub-Saharan Africa [*33,34*] and provide protection against African sleeping sickness, which is endemic to sub-Saharan Africa. Therefore, it has been assumed that they are prevalent in these populations due to their heterozygote advantage alone, much as in the case of the HbS mutation with regard to malaria protection (*26, 27, 29, 30*). Thus, examination of the de novo origination rate of the mutations involved can provide empirical test cases similar to that of the HbS mutation in distinguishing whether the prevalence of an adaptive mutation in a population is due to random mutation and natural selection (plus subsidiary factors such as random genetic drift) or whether such mutations are more likely to arise de novo in the populations where they are adaptive.

Melamed et al.’s method of Mutation Enrichment followed by upscaled Maximum Depth Sequencing (MEMDS) has enabled measurement of mutation rates of individual mutations by focusing on narrow genomic regions of interest (ROIs) and combining high throughput consensus sequencing with a mutation enrichment step (*12*). A key element of it is the controlled removal of the vast majority of wild-type (WT) gene-copy fragments prior to sequencing using a restriction enzyme (RE) that digests specifically WT molecules while leaving RE-protected mutant sequences intact (*12*). This removal of WT greatly reduces the number of WT molecules that undergo sequencing and which could have become false positive mutation calls. The molecules removed are known to carry the WT sequence in the ROI and therefore need not be sequenced. This step reduces both sequencing runs and the false positive rate due to PCR amplification or high-throughput sequencing errors by the same factor. Next, randomized barcodes are attached directly to the remaining ROI-carrying molecules, allowing linearly amplified copies that originate from the same DNA fragment to be grouped together and consensus sequenced to remove the remaining PCR and high-throughput sequencing errors at the computational stage (*35*). At the same time, the novel experimental design described in ref. (*12*) is used to keep track of the number of WT molecules removed and, based on it, calculate the frequencies of mutations in the original sample. For this purpose, two DNA samples are prepared, each with the addition of mock DNA resistant to RE digestion, and both are subjected to the same protocol, except for the RE digestion step, which is applied to only one of them. Following accurate barcode-based consensus sequencing, one obtains the ratios of mock RE-resistant to RE-sensitive sequences in each sample, and, based on them, calculates the enrichment factor of RE-resistant mutations, the number of WT molecules removed by digestion, and thus the frequency in the original sample of each RE-resistant mutation (Figure 1A). This method lowers the sequencing runs and error rate sufficiently to identify and count de novo mutations in human sperm samples at the single-digit resolution (*12*).

**Figure 1:**
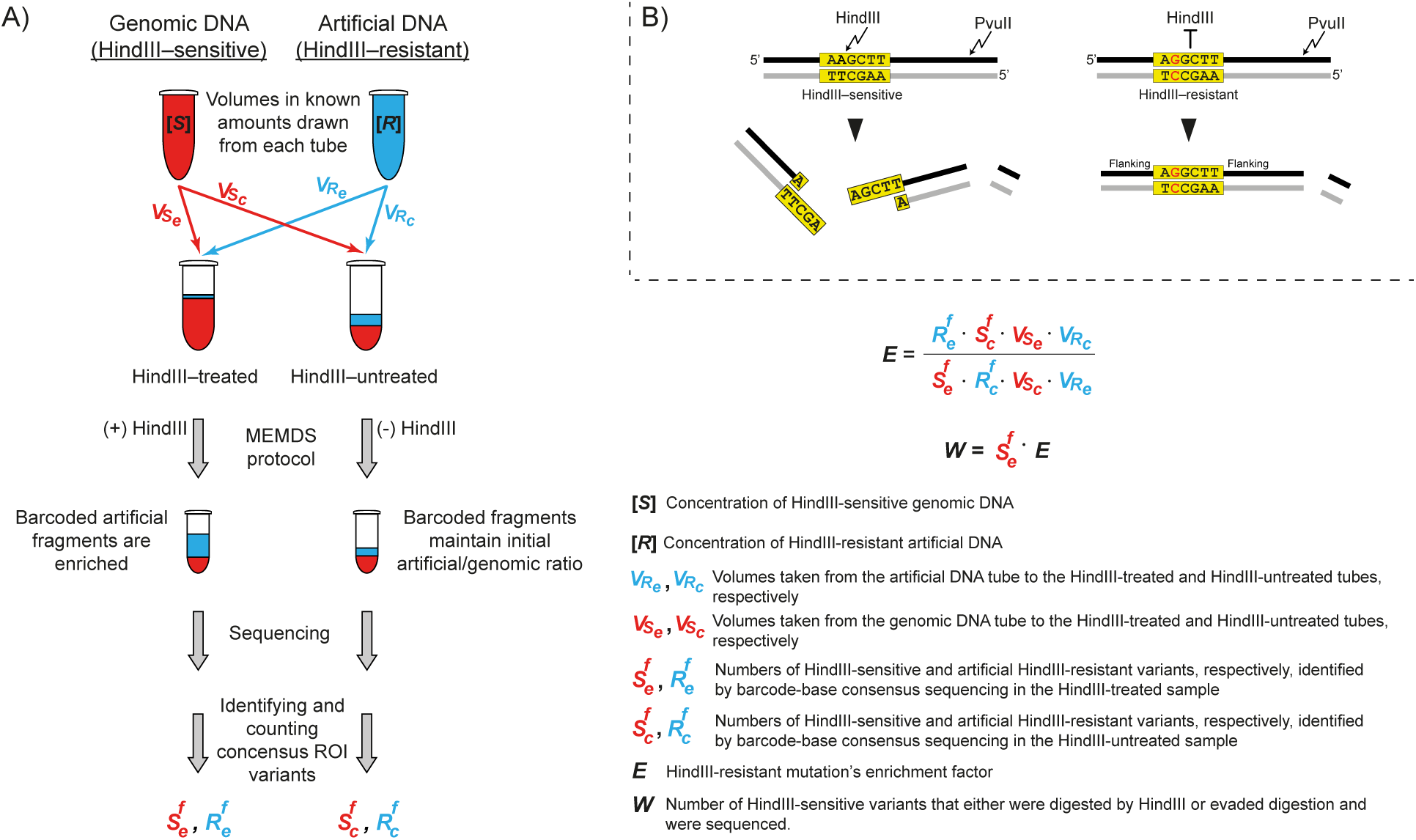
MEMDS experimental setup. A) Schematic illustration of the MEMDS experimental design enabling calculation of the HindIII-enrichment factor and the number of target-*APOL1* WT molecules that were scanned, modified from ref. (*12*). The experiment begins with two tubes, one containing sperm genomic DNA that carries mostly HindIII–sensitive *APOL1* ROI sequences, *S*, and one containing artificial *APOL1* molecules, *R*, which carry a known 6 bp sequence different from the HindIII recognition sequence and are therefore resistant to HindIII digestion. Volumes are drawn in known amounts from each source tube to create two mixtures designated “HindIII–treated” and “HindIII–untreated.” These two samples undergo the full MEMDS protocol (Figure S1), with the exception that the former is treated with HindIII to enrich for mutations in the ROI while the latter is not. Note that the volumes taken for both mixtures are not equal to allow for efficient identification of both genomic and artificial *APOL1* sequences following the different treatments and to scan a large amount of genomic *APOL1* sequences for de novo mutations in the HindIII-treated sample (detailed in ref. (*12*), Supplementary Information Text 2). Following highthroughput sequencing, variants are identified by the MEMDS computational pipeline (*12*), and the numbers of HindIII-sensitive ROI variants (i.e., WT ROIs) and artificial RE-1–resistant ROI variants are determined for each sample. These numbers, together with the known volumes taken from the source tubes to create the input mixtures, are used to calculate the HindIII enrichment factor, *E*, and the total number of WT sequences that were either digested by HindIII or evaded digestion and were sequenced and identified, *W*, as shown in the figure (for full derivation and assumptions, see ref. (*12*)). B) A schematic illustration of the *APOL1* gene fragment analyzed and variant enrichment. The 6 bp region of interest (ROI) that matches the HindIII-restriction site is shown in the yellow box. HindIII digestion of human sperm DNA removes *APOL1* gene fragments carrying the WT sequence and enriches for *APOL1* variants with mutations at the ROI site, while the experimental setup described enables keeping track of the number of WT molecules removed. PvuII digestion is used for the addition of a unique barcode sequence near the ROI for the subsequent consensus sequencing. See Figure S1 for the complete MEMDS protocol.

In the case of mutations that are expected to affect the fitness of the organisms carrying them but not the viability or fertility of sperm cells of healthy non-carriers in which they appeared de novo, such as the HbS or the *APOL1* G1 or G2 mutations, the frequency of the mutation in sperm samples is equivalent to the probability that it will be transmitted to the offspring and appear as a de novo mutation in it. Therefore, it is equivalent to the evolutionarily relevant mutation-specific de novo origination rate. Importantly, this equivalence holds regardless of whether different instances of such a mutation in the sample have all arisen independently or via replications of a cell or cells that mutated early in spermatogenesis, a possibility referred to in ref. (*12*) as “clonal dependence.” Although clonal dependence, if it exists in the data, would increase the variance in the mutation rate estimate, this has been taken into account statistically both in ref. (*12*) and here.

For its WT-reduction procedure, MEMDS requires both the ROI to consist of an RE recognition sequence and the mutations of interest to disrupt this sequence. Therefore, we examined the G1 and G2 mutations and found that the G1 1024A*→*G mutation satisfies these conditions: it disrupts a sequence that, in the WT, constitutes the recognition site of HindIII (AAGCTT), a commercially available RE (Figure 1B). Accordingly, we obtained sperm samples from 7 Ghanaian and 8 northern European donors and examined a total of more than half a billion *APOL1* ROI-carrying gene-copy fragments individually to measure the de novo origination rates of the G1 1024A*→*G mutation as well as nearby point mutations and indels in or overlapping with the *APOL1* ROI. See Supplementary Information for a complete methods description and sequence library properties (Figures S1–S10 and Tables S1–S4).

## Results

Previous studies utilizing single-strand DNA sequencing consider G*→*T and C*→*T changes to represent the experimental disruption of an ongoing in vivo process of base damage and repair due to guanine oxidation and cytosine deamination as well as corresponding *in vitro* damage rather than durable mutations which might be propagated (*12, 35, 36*). Indeed, in accord with this, the vast majority of DNA sequencing variations observed here were of these two types (Figure S6) and were excluded from further analysis.

In addition, no natural de novo mutations are expected to be observed in notable frequencies in the ROI in the HindIII-untreated samples, and, if they are observed there, they should appear at incomparably lower frequencies compared to the HindIII-treated sample due to lack of enrichment. However, three mutations, 1026C*→*A, 1027T*→*A and 1028T*→*A were observed in the ROI in the untreated samples (no other mutations, including the G1 1024A*→*G mutation, were observed in these samples), and their frequencies, normalized to the average noise level in the ROI-flanking regions, were very similar between the treated and untreated samples (Figure S10). These observations are inconsistent with the possibility that these mutations originated naturally during gametogenesis and were enriched with MEMDS, but are consistent with the possibility that they represent noise introduced during library preparation. Therefore, we excluded these mutations from further analysis. Following these foregoing exclusions, 13 point mutations and potentially many indels remain observable by MEMDS in this ROI (Table 1).

**Table 1:** Counts of de novo mutations observable by MEMDS in the *APOL1* ROI in sperm samples, 7 from sub-Saharan Africans and 8 from northern Europeans. The number of haploid individual genomes scanned by MEMDS in the ROI per donor are given in the second column in units of 10^6^. The base positions of observable mutations and the particular observable mutations in those positions are given in the second and third rows for both point mutations and indels (all 4 observed indels are single-base deletions). The other rubrics provide the counts of de novo instances per donor for each mutation (empty cells represent zero counts). The rate of the 1024A*→*G mutation is notably higher in Africans than Europeans and notably higher than that of all other observable mutations in the region.

| Positions (or center of indels)→ |  | Point mutations |  |  |  |  |  |  |  |  |  |  |  | Indels |  |  |  |  |
| --- | --- | --- | --- | --- | --- | --- | --- | --- | --- | --- | --- | --- | --- | --- | --- | --- | --- | --- |
|  |  | 1023 |  |  | 1024 |  |  | 1025 |  | 1026 | 1027 |  | 1028 |  | 1024 del | 1025 del | 1026 del | 1028 del |
| Cells scanned (10<sup>6</sup>)↓ |  | A>G | A>C | A>T | A>G | A>C | A>T | G>A | G>C | C>G | T>C | T>G | T>C | T>G |  |  |  |  |
| AFR | 54.0 | 2 |  |  | 9 |  |  |  |  |  | 1 |  |  |  |  |  |  |  |
|  | 39.4 |  |  |  | 5 |  |  |  |  | 1 | 1 |  | 1 |  |  |  |  |  |
|  | 50.3 | 2 | 1 |  | 1 |  |  | 1 | 1 |  | 2 |  | 1 |  |  | 1 |  | 1 |
|  | 33.1 | 4 |  |  | 4 |  |  |  |  | 1 | 1 |  | 3 |  |  |  |  |  |
|  | 38.5 | 1 |  |  | 11 |  |  |  | 2 |  |  |  | 2 |  |  |  |  |  |
|  | 30.8 | 4 |  |  | 7 |  |  |  |  | 1 |  |  | 5 |  |  |  |  | 2 |
|  | 45.0 | 3 |  |  | 12 |  |  | 7 |  |  |  |  |  |  | 1 |  |  |  |
| EUR | 47.4 | 1 |  |  |  |  |  |  |  | 1 |  |  |  |  |  |  |  |  |
|  | 54.9 |  |  |  | 2 |  |  |  |  |  |  |  | 1 |  |  |  |  |  |
|  | 43.7 |  |  |  | 4 |  |  | 1 |  |  | 2 | 1 | 3 |  |  |  | 1 | 1 |
|  | 36.8 | 2 |  |  | 5 |  |  |  |  | 2 |  |  | 1 |  | 1 |  |  |  |
|  | 20.2 |  |  |  |  |  |  |  |  |  |  |  |  |  |  |  |  |  |
|  | 45.8 | 2 |  |  | 3 |  |  |  |  |  |  |  | 2 |  |  |  |  |  |
|  | 35.3 | 1 |  | 1 | 7 |  |  |  |  |  | 3 |  | 3 | 1 |  |  |  |  |
|  | 38.6 | 1 |  |  |  |  |  |  | 2 |  | 1 |  |  |  |  | 1 |  |  |

The mutations observed in the flanking regions outside of the ROI are not enriched. Although some of these mutations may be real, categorizing all of them as due to noise and dividing the average mutation rate across the flanking regions by the enrichment factor sets an upper bound on the noise level (the false positive rate, FPR) in the ROI (Figure 2). While in ref. (*12*) this upper bound FPR was itself negligible (10*^−^*^9^), for the current ROI it was found to be on the order of 10*^−^*^8^. Although in principle this noise level could make a study of mutation rates at the single mutation resolution challenging given the average human genome-wide mutation rate of *∼*10*^−^*^8^, here it turned out not to be the case because of the hotness observed for this ROI. Nevertheless, in order to take this noise level into account and obtain the lower bound (i.e., most conservative) de novo mutation rates, we subtracted the upper bound FPR of each mutation from the actual observed rate of that mutation calculated from the numbers in Table 1, obtaining the mutation rates in Figure 3. All statistical tests and their results provided below are based on these noise-reduced numbers, and in all count-based tests the expected mutation numbers following noise reduction were first summed up and then rounded to the nearest integer. Following this upper bound noise reduction, the total numbers of point mutations in Africans and Europeans were 88 and 47 respectively (compared to 97 and 53 in Table 1), showing that even this maximum possible noise reduction has a limited effect on the data.

**Figure 2:**
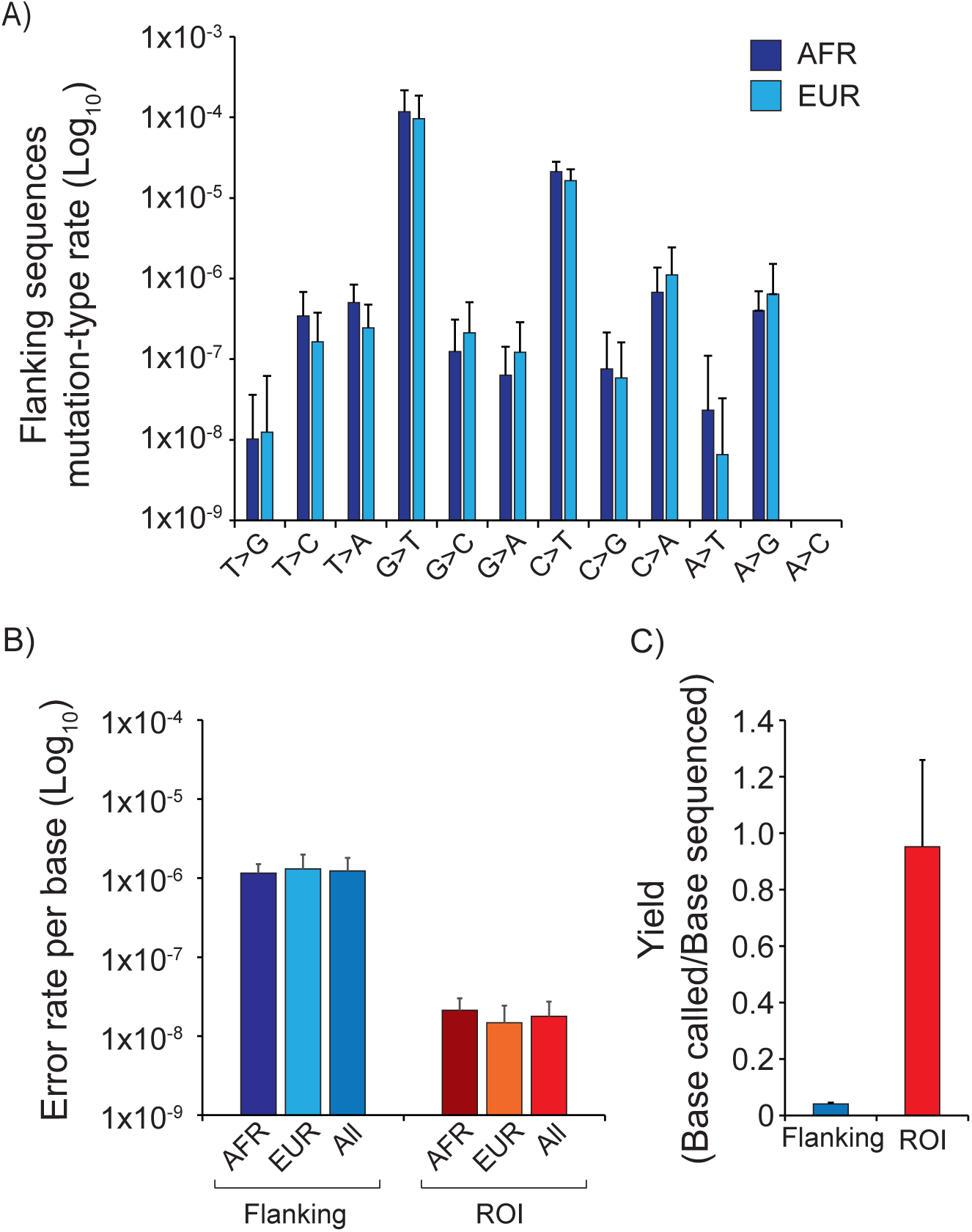
Error rates and yield of MEMDS as applied to the *APOL1* ROI. A) Rates per mutation type in the ROI-flanking sequences of *APOL1* fragments for the HindIII-treated and HindIII-untreated samples combined show a similar pattern for the African (AFR, dark blue) and European (EUR, light blue) groups. B) Under the stringent assumption that all of the mutations in the flanking regions are due to sequencing and/or library generation errors, these mutations, with the exception of G*→*T and C*→*T, are used to calculate the average FPR per base in the flanking sequences (blue). To calculate the average FPR for the 6 bp ROI (red), the per-donor FPR rates for the flanking sequences are divided by the corresponding HindIII-enrichment factors. C) Depletion of *APOL1* sequences carrying the WT ROI by HindIII digestion increases the yield (i.e., the number of called consensus bases per sequenced base) for the 6 bp ROI sequence. The yield can become larger than one because the composition of the sequences removed by digestion is known.

**Figure 3:**
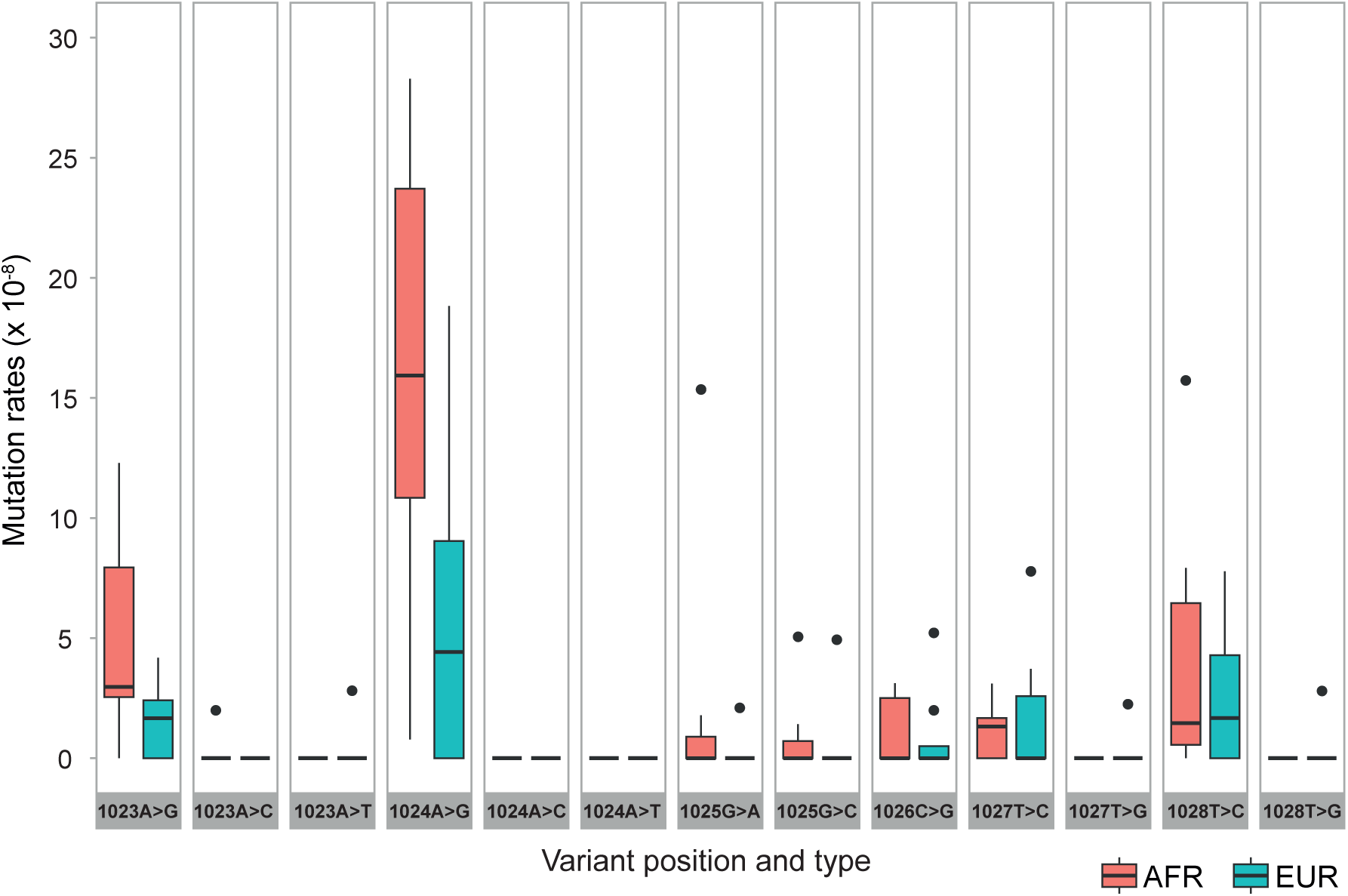
Boxplot of de novo mutation rates of observable point mutations in the *APOL1* ROI in Africans (AFR, red) and Europeans (EUR, turquoise). The *Trypanosoma brucei gambiense*–protective 1024A*→*G mutation exhibits a striking difference in de novo rate between the two populations.

### Average ROI mutation rates

To obtain comparisons to well-accepted quantities, we used 1.25 *×* 10*^−^*^8^ as grossly representative of the human genome-wide average point mutation rate [*2, 3, 37–40*] and a 10*×* lower estimate (1.25 *×* 10*^−^*^9^) for the human genome-wide average indel rate (*8, 12, 41, 42*). We found that, as measured by MEMDS for the two ethnic groups combined, the average point mutation and indel rates in the *APOL1* ROI were 5.1 *×* 10*^−^*^8^ and 2.4 *×* 10*^−^*^9^ respectively, significantly higher by *∼*4.1-fold (*P <* 5 *×* 10*^−^*^40^, 95% CI 4.3 *×* 10*^−^*^8^–6.0 *×* 10*^−^*^8^, two-sided binomial exact test) and marginally significantly higher by *∼*2.0-fold (*P* = 0.055, 95% CI 1.1 *×* 10*^−^*^9^–4.6 *×* 10*^−^*^9^, two-sided binomial exact test) than the aforementioned genome-wide average point mutation and indel rates respectively.

### Population differences in ROI rates

To test for a population-level difference in the overall point mutation rates per person across the *APOL1* ROI while excluding the possibility that such a difference is merely due to sample-level variation (e.g., due to clonal dependence) and/or individual-level variation alone, we used a two-sided Wilcoxon rank-sum test. We found that these rates were marginally significantly higher in the African than in the European group (*P* = 0.054). When excluding the G1 1024A*→*G mutation from the count, the rates were not significantly different between the groups (*P* = 0.15, two-sided Wilcoxon rank-sum test). Next, pooling together cells from all donors within each group while ignoring individual- and sample-level variation to obtain the average point mutation rate including the G1 1024A*→*G mutation across the ROI showed this average rate to be significantly higher by *∼*2.1-fold (*P <* 10*^−^*^4^, OR 95% CI 1.45–3.00, two-sided Fisher exact test) in Africans than in Europeans. The average pooled indel rate did not differ significantly between the groups (*P* = 0.74, two-sided Fisher exact test).

### Mutation-specific origination rates

While the above measurements relate to population comparisons for the overall ROI, statistical analysis of the results at a finer scale resolution yielded a clear picture. First, following upper-bound noise reduction, analysis showed that nearly 46 G1 1024A*→*G mutation instances were observed in the African group among a total of *∼*291 *×* 10^6^ cells, whereas 19 G1 1024A*→*G mutation instances were observed in the European group among a total of *∼*323 *×* 10^6^ cells. To examine whether the de novo origination rates of the G1 1024A*→*G mutation differed significantly between the groups while excluding the possibility that this difference is due to sample-level variation (e.g., due to clonal dependence) and/or individual-level variation alone, we compared the per person G1 1024A*→*G mutation rates between the two groups using a two-sided Wilcoxon rank-sum test. Results showed that these rates were significantly higher in the African than in the European group (*P* = 0.039). Next, pooling together cells from all donors within each group while ignoring individual- and sample-level variation to obtain the average mutation-specific origination rate of G1 1024A*→*G across individuals showed this rate to be significantly higher by *∼*2.7-fold (*P <* 2 *×* 10*^−^*^4^, OR 95% CI 1.56–4.72, two-sided Fisher exact test) in Africans than in Europeans.

Additionally, we found that the G1 1024A*→*G mutation is the point mutation of highest mutation-specific rate in the ROI (Figure 3), and that it deviates by far the most from the genome-wide average mutation rate for its type (A*→*G) compared to all other 12 point mutations studied in the ROI (Table 2). This deviation increases when expanding the local genetic context (the size of the motif) based on which the genome-wide average mutation rates are calculated, from 1-mer to 7-mer (Table 2).

**Table 2:** de novo mutation rate fold changes from the corresponding genome-wide average (GWA) rates for observable point mutations in the African *APOL1* ROI. GWA rates were calculated for the 1-mer context based on genome-wide family sequencing studies (*3*) and for the 3-, 5- and 7-mer contexts based on the relative frequencies of Extremely Rare Variants (ERVs) (*5*). The mutation-specific origination rates of the 1024A*→*T mutation deviates by far the most from the GWA for its type compared to all other 12 observable point mutations in the ROI (over 26-fold vs. *∼*8- and *∼*6-fold for the second and third largest deviations in the 1-mer context, and nearly 57-fold compared to 11- and 15-fold in the 7-mer context). The deviations are not accounted for by expanding the local genetic context taken into account in the GWAs. Adjustments to the GWA calculation are minor compared to the variation in mutation-specific origination rates, and the deviations of the three mutations of highest rate from their corresponding GWAs merely increase overall as we move from the 1-mer to the 7-mer context.

|  | *de novo* mutation based |  | ERV-based, 3-mer |  | ERV-based, 5-mer |  | ERV-based, 7-mer |  |
| --- | --- | --- | --- | --- | --- | --- | --- | --- |
| Variant | GWA rate (x10<sup>-8</sup>) | Fold-change | GWA rate (x10<sup>-8</sup>) | Fold-change | GWA rate (x10<sup>-8</sup>) | Fold-change | GWA rate (x10<sup>-8</sup>) | Fold-change |
| 1023A>G | 0.593 | 8.11**** | 0.452 | 10.65**** | 0.531 | 9.05**** | 0.443 | 10.85**** |
| 1023A>C | 0.170 | 2.03 | 0.146 | 2.35 | 0.121 | 2.84 | 0.125 | 2.74 |
| 1023A>T† | 0.138 | 0.00 | 0.176 | 0.00 | 0.108 | 0.00 | 0.116 | 0.00 |
| **1024A>G** | 0.593 | 26.64***** | 0.431 | 36.64***** | 0.314 | 50.37***** | 0.278 | 56.93***** |
| 1024A>C† | 0.170 | 0.00 | 0.157 | 0.00 | 0.150 | 0.00 | 0.125 | 0.00 |
| 1024A>T† | 0.138 | 0.00 | 0.105 | 0.00 | 0.099 | 0.00 | 0.079 | 0.00 |
| 1025G>A | 0.732 | 3.76** | 0.829 | 3.32** | 0.802 | 3.43** | 0.840 | 3.27** |
| 1025G>C | 0.213 | 4.83* | 0.238 | 4.34* | 0.300 | 3.44 | 0.264 | 3.90* |
| 1026C>G | 0.213 | 4.83* | 0.238 | 4.34* | 0.300 | 3.44 | 0.206 | 5.00* |
| 1027T>C | 0.593 | 1.74 | 0.431 | 2.39 | 0.402 | 2.56 | 0.353 | 2.92 |
| 1027T>G† | 0.170 | 0.00 | 0.157 | 0.00 | 0.147 | 0.00 | 0.184 | 0.00 |
| 1028T>C | 0.593 | 6.37*** | 0.322 | 11.74**** | 0.329 | 11.50**** | 0.252 | 15.01**** |
| 1028T>G† | 0.170 | 0.00 | 0.135 | 0.00 | 0.155 | 0.00 | 0.741 | 0.00 |

† Variant was not found in the analyzed samples of African donors
\* $p < 5 \times 10^{-2}$
\*\* $p < 5 \times 10^{-3}$
\*\*\* $p = 2.11 \times 10^{-6}$
\*\*\*\* $p < 7 \times 10^{-9}$
\*\*\*\*\* $p < 3 \times 10^{-48}$

We also found that the lower frequency 1023A*→*G mutation shows a significant difference in de novo rates between the populations (*P* = 0.042, two-tailed Wilcoxon rank-sum test), albeit with much smaller rates.

## Discussion

Results showed that the G1 1024A*→*G mutation, which protects against *Trypanosoma brucei gambiense*—the most common cause of human African sleeping sickness, originated de novo significantly more frequently in the African than in the European group. Namely, the G1 1024A*→*G de novo rate is significantly higher in the population where this mutation is of adaptive significance. Furthermore, the pattern held even when controlling for possible clonal dependence among the observed instances of the mutation. Adding to the uniqueness of the G1 1024A*→*G mutation, not only is its rate by far the highest among the rates of all other point mutations in the region, it also deviates by far the most from the genome-wide average for its type (A*→*G), and this deviation merely increases when expanding the local genetic context (further highlighting the fact that the local genetic context alone does not explain mutation rate variation at the single mutation resolution).

These results show a strikingly similar pattern to that recently observed for the malaria-protective HbS mutation in the African *HBB* gene (*12*). In that study, Melamed et al. found that the de novo HbS mutation origination rate was higher specifically in *HBB* in Africans—i.e., in the gene and population case where it protects against malaria—compared to the other three groups combined (the *HBB* gene in individuals of European origin, whose ancestors have not been exposed to intense malarial pressure, and the paralogous *HBD* gene, which is not implicated in malaria resistance, in individuals of both African and European origin), while accounting for possible clonal dependence (*12*). In addition, they addressed the question of whether the effect existed at the population level independently of a gene-level effect as follows. They noted that, both with or without counting the HbS mutation, the overall point mutation rate in the *HBB* ROI was significantly higher in Africans than in Europeans and a significant correspondence existed between the de novo mutation rates observed in the study and allele frequencies in populations (*12*); and that for the HbS mutation rate not to be higher in Africans than in Europeans, it would have had to violate both of these independent statistically significant patterns, when in fact it was already aligned with both more strongly than the rate of any other point mutation in the region (*12, 43*). In other words, triangulating from different statistically significant tests suggested that the population-level difference held independently of a gene level one [*12,43*]. In contrast, in the current study, the higher mutation rate for the *APOL1* ROI compared to the *HBB* ROI together with a larger sample size yielded a larger number of observations sufficient to resolve with a single, simple statistical test that the G1 1024A*→*G mutation rate is higher in Africans than in Europeans.

It is thus notable that in both of the first two studies where it has been possible to examine mutation-specific de novo origination rates, both of which examined mutations of known adaptive values—the well-known malaria-protective HbS mutation as well as the G1 1024A*→*G mutation— the focal mutations originated de novo significantly more frequently specifically where they were of adaptive value. Both cases thus add a level of complexity to what was previously known regarding mutation rates (e.g., refs. [*3, 5, 11, 37, 38, 44*]). Indeed, while it has been assumed that “the likelihood of any particular mutational event is independent of its specific value to the organism” (*20*), both studies showed a pattern opposite than expected. It is worthwhile to note that no studies of mutation rates at the single mutation resolution existed before. All past conclusions on mutation rates had been based on mutation rate averages across many genomic positions [*3–8, 45*]. Therefore, unless the new findings are refuted experimentally, they require a reexamination of the aforementioned expectation and its consequences (*46*).

Melamed et al. offered several possible explanations for the HbS mutation findings, which we can consider together with the current convergent pattern shown for the *APOL1* G1 1024A*→*G mutation (*12*). First, it could be a mere coincidence of population-dependent, locus-specific genomic fragility that the G1 1024A*→*G mutation originates more frequently in *APOL1* in sub-Saharan Africans, that it protects against African sleeping sickness, and that the latter is endemic in sub-Saharan Africa; and likewise that due to another population-dependent, locus-specific genomic fragility, the HbS mutation originates more frequently in sub-Saharan Africans and protects against malaria, which is endemic in sub-Saharan Africa. However, this interpretation would leave anomalous data from both of the first two consecutive studies on mutation specific origination rates, as in fact the results from both are inconsistent with the textbook definition of random mutation (*21*) (see also ref. (*20*)).

A second possibility is that the adaptive mutation rates can be attributed to modifier alleles which themselves originated via random mutation in the past and were consequently selected due to their beneficial effects on the mutation rate. However, this formulation seems untenable in light of the fact that DNA mutation rates are low. In order for a modifier allele to be selected based on its effect on the mutation rate, it would need to affect the average mutation rate across many genomic positions, so that many potential mutation events could figure into its selective benefit [*11, 47–50*]. Thus, modifier theory in its present form cannot explain the sharp deviations of individual DNA mutations in specific base positions from their corresponding genome-wide average rates observed in the two studies.

A third possibility is that mutation-specific origination rates depend on complex information accumulated in the genome over generations [*12, 46, 51–54*]. For example, it has been observed recently that genes that are used together repeatedly over generations are more likely to become fused together by mutational mechanisms based on their interactions (*52, 53*); that positions that are repeatedly edited over generations at the RNA level may be more likely to undergo the corresponding change at the DNA level due to mutational mechanisms based on RNA editing (*53*), involving RNA editing in the mechanisms of evolution (*55*); etc. (*53*). Although this possibility leaves open for future studies to establish the precise mechanisms that generate the high HbS and G1 1024A*→*G de novo mutation rates, the overall principle here is that mutation-specific origination rates are influenced by preexisting genetic interactions [*12, 46, 51–54*]. Some observations could be further brought to bear on the problem, such as that strong A-to-I RNA editing activity (effectively generating A*→*G change) has been described in the 3’UTR of *APOL1*, which is relatively close to the *APOL1* ROI, although no such activity was found in the 1024A*→*G site itself under the studied conditions (*56–58*). Accordingly, the generation of heritable change could be neither random nor Lamarckian: it could be influenced by biologically meaningful information that does not necessarily arrive directly from the immediate environment but is accumulated internally in the genome over generations, in a manner specific to each heritable change [*12,46,51–54*]. It will be important to test this possibility further by using ultra-high accuracy mutation detection and other methods in a variety of genes, organisms and populations and across mutation types.

## Materials and Methods

The MEMDS method as applied to the *APOL1* G1 mutation was adapted from Melamed et al. (*12*). A detailed description of the MEMDS method is available in the Supplemental Text and Figure S1.

### Reagents

All oligos were obtained from Integrated DNA Technologies (IDT) with standard desalting purity. Full descriptions of the oligos used are available in Table S4. All enzymes were received from New England Biolabs (NEB). DNA purifications were carried out using QIAGEN kits in accord with the manufacturer’s instructions unless mentioned otherwise.

### Preparation of spike-in plasmids

Two puc19-based plasmids, ALP23 and ALP24, were constructed to include a sequence identical to the studied *APOL1* gene fragment between positions 764 and 1255, relative to the mRNA translation start site, except that the HindIII AAGCTT restriction site was replaced with TTGAAA and CTCTAG, respectively. To make the spike-in mixture, the two plasmids were linearized by BamHI, mixed in equal amounts and diluted to 10 fg/µl.

### Collection of sperm samples

Sperm samples were obtained from healthy donors between the ages 18 and 39 with no history of cancer or infertility and with no high fever in the 3 months prior to donation. European samples were purchased from Fairfax Cryobank and African samples were collected in the Assisted Conception Unit of the Lister Hospital & Fertility Centre in Accra, Ghana following clinical standards with the approvals of the Institutional Review Board of the Noguchi Memorial Institute for Medical Research (NMIMR-IRB 081/16-17) at the University of Ghana, Legon, the Rambam Health Care Center Helsinki Committee, Haifa (0312-16-RMB) and the Israel Ministry of Health (20188768). Informed consent was obtained from all participants, and personal identifying information was removed and replaced with codes at the source.

### Extraction of Sperm DNA

DNA extraction from sperm samples was carried out as described previously by Melamed et al. (*12*). Briefly, 500 µl aliquots from each donor’s sperm sample were washed twice with 70% ethanol to remove seminal plasma. Cells were rotated overnight at 50°C in a 700µl lysis buffer (50 mM Tris-HCl pH 8.0, 100 mM NaCl2, 50 mM EDTA, 1% SDS) containing 0.5% Triton X-100 (Fisher BioReagents BP151-100), 50mM Tris (2-carboxyethyl) phosphine hydrochloride (TCEP; Sigma-Aldrich 646547) and 1.75 mg/mL Proteinase K (Fisher BioReagents BP1700-100). Lysates were centrifuged at 21,000×g for 10 minutes at room temperature and supernatants were united in a single tube. The cleared lysate was supplemented by 15 mL of buffer G2 (QIAGEN Blood & Cell Culture DNA Maxi Kit, 13362), and purification continued according to the manufacturer’s instructions. The eluted DNA was allowed to dissolve overnight at room temperature. For each donor, a small aliquot from the extracted DNA was PCR-amplified and sanger-sequenced to confirm the absence of G1 and G2 mutations in each donor’s genetic background as well as to confirm the compatibility of the target region for library generation by the MEMDS protocol (i.e., having no other somatic mutations that may inhibit HindIII or PvuII digestion or which may interfere with primer binding). Additionally, the haplotype background at positions 150, 228 and 255 of the *APOL1* protein were verified for all donors.

### Enzymatic digestion of Sperm DNA

For the HindIII-treated sample, DNA amounts equivalent to 75-90 million haploid cells were mixed with 0.2 pg of the plasmid spike-in mixture, divided in a 96-well plate with 10 U HindIII-HF (R3104L) per well and incubated overnight at 37°C, according to the manufacturer’s instructions (for the genomic and plasmid spike-in mixture volumes used that were crucial for the HindIII enrichment factor calculation, see Table S3). Then, each well was supplemented by 20 U PvuII-HF (R3151L) and 20 U NdeI (R0111L) and incubated for an additional 3 hours at 37°C.

For the HindIII–untreated reaction, sperm DNA in the amount equivalent to 5% of that used for the HindIII-treated reaction was mixed with 1.2 pg of plasmid spike-in mixture and 10 units SalI-HF (R3138L), aliquoted to 5 tubes, and incubated overnight at 37°C, according to the manufacturer’s instructions. This allowed for similar conditions of DNA digestion, without affecting *APOL1* target sequences. Each tube was then supplemented by 20 U PvuII-HF (R3151L) and 20 U NdeI (R0111L) and incubated at 37°C for an additional 3 hours. DNA purification was carried out with QIAGEN PCR purification kit.

### Primary barcode labeling and linear amplification

Direct barcode labeling followed by linear amplification of the digested *APOL1* strands were carried out together in a single reaction in 96-well plates. Each well contained about 1µg of digested DNA, 0.1 µM primary barcode oligo (Oligo APA; see Table S4) and 1 µM of 5’-phosphorothioate-protected primer for linear amplification (Oligo APB). The reaction was carried out with Q5 high-fidelity polymerase according to the manufacturer’s instructions, using the following thermocycler parameters: initial denaturation at 98°C for 20 seconds, followed by 16 cycles of 98°C for 5 seconds, 67°C for 15 seconds, and 72°C for 20 seconds. For each donor and each treatment, a different APA oligo was used with a unique Donor Identifier-1 (ID-1) sequence, allowing for elimination of any post-processing contamination at the sequence analysis step.

### 5’-exonuclease treatment

Following purification, ∼15 µg DNA aliquots from the post–linearly amplified product of the HindIII-treated sample were incubated, each at 37°C, in the presence of 20 U of Lambda exonuclease, 40 U of T7 exonuclease and 120 U of RecJF exonuclease in 1x CutSmart buffer for 3.5 hours. The post-linearly amplified product of the HindIII-untreated sample was incubated at 37°C for 3.5 hours with 10 U of Lambda exonuclease, 20 U of T7 exonuclease and 60 U of RecJF exonuclease.

### Secondary barcode labeling and 3’-exonuclease treatment

Following purification, the DNA was aliquoted into a 96-well plate (∼0.75 µg per well). A single primer extension reaction was carried out using 0.5 µM of the secondary barcode primer (Oligo APC) and Q5 high-fidelity polymerase according to manufacturer’s instructions, using the following thermocycler parameters: initial denaturation at 98°C for 20 seconds, followed by a single cycle of 98°C for 5 seconds, 70°C for 15 seconds, and 72°C for 40 seconds. Immediately after the thermocycler temperature dropped to 16°C, to remove free oligo APC, 20 units of thermolabile Exo I were added directly to each well, together with the relabeling control primer (Oligo APD). Oligo APD was added in a known amount equivalent to 0.66% of the secondary barcode primer (6.6% for AFR3 and AFR7). After one hour of incubation at 37°C, Exo I was heat-inactivated for one minute at 80°C and the DNA was purified. For each donor, each of the HindIII-treated and untreated samples was labeled by an Oligo APC with a different Donor Identifier-2 sequence (ID-2), which also was not shared by samples from other donors, resulting in each donor and each condition having a unique Identifier-2 sequence.

### PCR amplification and sequencing

The first PCR reaction of the barcoded product was carried out using the primers APE and APF1 and Q5 high-fidelity polymerase according to manufacturer’s instructions. The following thermocycler parameters were used: initial denaturation at 98°C for 30 seconds, followed by 10 cycles of 98°C for 5 seconds, 72°C for 15 seconds, 72°C for 30 seconds, and a final extension at 72°C for 30 seconds. Amplification products were purified and the second PCR reaction was carried out using 25% of the first PCR product as a template, the primers APE and APF2, and Q5 high-fidelity polymerase according to the manufacturer’s instructions (different F2 primers were used in order to add a unique Illumina index sequence to each HindIII-treated and untreated sample). The following thermocycler parameters were used: initial denaturation at 98°C for 30 seconds, followed by 20 cycles (for AFR1-4 and EUR1-4) or 16 cycles (for AFR5-7 and EUR5-7) of 98°C for 5 seconds, 70°C for 15 seconds, 72°C for 30 seconds, and a final extension at 72°C for 1 minute. PCR products were agarose-gel–purified and further concentrated by a DNA clean & concentrator kit (Zymo Research). DNA libraries prepared from the HindIII-treated and untreated samples of the same donor were mixed in a 3:2 ratio, respectively. HindIII-treated and untreated mixtures from three donors were combined and paired-end sequenced with 50% control yeast-genomic DNA library by Illumina NextSeq 550 Mid Output 300 cycles at the Technion Genome Center (TGC) and at the Genomic Center of the Azrieli Faculty of Medicine, Bar-Ilan University.

### Sequence analysis

Paired-end (PE) sequences were merged via Pear (*59*) using the default options. Merged sequences were trimmed from Illumina adapters using Cutadapt (*60*) and quality-filtered by Trimmomatic (*61*), using a sliding window size of 3 bp, a Phred quality threshold of 30 and a minimum read length threshold of 90 bp. Quality filtered sequences were trimmed at their edges to retrieve barcode and sample ID data from each read. The 5’ edge was trimmed up to position 18, to include the 14 bases of the primary barcode and the 4 bases of sample ID-1. Similarly, the last 9 bp at the 3’ edge was trimmed to include the 5 bases of the secondary barcode and the 4 bases of sample ID-2. The barcode and the ID information was added to the read header. Only sequences with the correct ID-1, ID-2 and first two bases of *APOL1* library sequence (consisting of the PvuII partial recognition sequence) were maintained. Trimmed sequences having the correct sample ID were confirmed for carrying the *APOL1* gene fragment based on the bases occupying positions 37-41 (ATTCG for *APOL1* sequence), allowing two mismatches and frameshifts of up to -3 or +3 bp upstream to the sorting positions. Approved sequences were mapped to the *APOL1* reference sequence (obtained by Sanger-sequencing aliquots from the matching donor samples) using BWA (*62*) with parameters -M -t. Query sequences whose first sequence position did not align with the first position of the reference sequence were excluded from further analyses. Mapped reads were scanned for presence of mutations using a custom script. High-quality mutations (Phred score *≥* 28) were noted. To confirm observed mutations as true variants, reads were grouped by their primary barcodes into “families” and processed according to the MEMDS pipeline (see Fig. S9 in Melamed et al. (*12*)). Briefly, only families containing at least four reads were included in the analysis; those not passing this cutoff were discarded. In the approved families, only mutations passing a mutation frequency cutoff of 0.7 and a secondary barcode cutoff of 3 were accepted. If a mutation failed to pass these thresholds, but the WT base in the same position was accepted by the same cutoff criteria, the base was considered WT. If neither mutation nor WT was accepted at a certain position, this position was considered ambiguous and marked by “N”. Read families having ambiguous positions were excluded from further analyses.

## Declarations

## Supporting information

SI text, tables and images

## Acknowledgements

We thank Mary Otoo and Joshua Adoboe for help with sample collection and Kim Weaver for extensive help. This publication was made possible through the support of grant 61129 from the John Templeton Foundation to AL and KS, ISF grant 3757/20 to KS, and grant 62220 from the John Templeton Foundation to AL (subaward). The opinions expressed in this publication are those of the authors and do not necessarily reflect the views of the John Templeton Foundation.

## Author contributions

DM and AL planned the experiments; MY and EH coordinated and carried out sample collection; DM, RS and DFB performed the experiments; EB and AL provided statistical tools; DM, EB and AL analyzed the data; MY, EH, KS and AL obtained Helsinki and IRB approvals; DM, EB and AL drafted the paper; DM, RS, EB, KS and AL revised the paper; KS provided general advice; KS and AL acquired funding; DM and AL conceived of the project; AL supervised the project.

## Declaration of interests

The authors declare that they have no competing financial interests.

## Declaration of AI use

The authors declare that AI assistance was not utilized during the analysis of experimental results and during manuscript writing.

## Supplementary information

Supplemental Text S1;

Supplemental Figures S1 to S10;

Supplemental Tables S1 to S4;

Supplemental References.

## Data and code availability

All raw sequencing data generated in this study is being submitted to the NCBI database of Genotypes and Phenotypes (dbGAP; https://www.ncbi.nlm.nih.gov/gap/) under accession number **phs002391.v2.p1**. For final processed data see Supplemental Datasheets and Supplemental Text.

The MEMDS pipeline is available at GitHub: https://github.com/livnat-lab/MEMDS_analysis_ *pipeline_v1.2*.

## Notes

### Competing Interest Statement

The authors have declared no competing interest.

