## Supplementary material for "De novo rates of a *Trypanosoma*-resistant mutation in two human populations": SI text, tables and images

**The SI file includes the below:**

Supplemental Text S1

Figures S1 to S10

Tables S1 to S4

Supplemental References

### Supplemental Text S1

#### 1 Applying the MEMDS method to study the *APOL1* 1024A→G mutation rate

##### 1.1 Method outline

###### 1.1.1 Enrichment and identification of de novo *APOL1* mutations by MEMDS

To study the de novo rates of the *APOL1* G1 (1024A→G) mutation and other, nearby mutations, we applied the Mutation Enrichment followed by upscaled Maximum-Depth Sequencing (MEMDS) method to human sperm DNA to enrich, sequence and identify mutations within a 6 bp region of interest (ROI) in the apolipoprotein L1 (*APOL1*) gene (Figure S1A). MEMDS, which was first applied to study the HbS mutation rate in the human *HBB* gene [*1*], utilizes two restriction enzymes: one to enrich for mutations at the ROI and the other to generate a priming site in order to add a unique primary barcode directly to the target DNA strand by a DNA polymerase-assisted fill-in reaction (Figure S1A). For the *APOL1* 1024A→G mutation enrichment we used the fact that the adenine base at position 1024 of the APOL1 coding sequence is a part of the HindIII restriction enzyme recognition site A<u>A</u>GCTT (position 1024 is underlined). Therefore, *APOL1* alleles carrying the WT sequence at the ROI (i.e., a complete HindIII site) are sensitive to HindIII digestion, whereas alleles carrying the 1024A→G mutation or any other mutation disrupting the HindIII site are resistant to digestion. Indeed, *APOL1* G1 resistance to HindIII digestion has long been used for genotyping purposes [*2*]. For the primary barcode addition, we used PvuII that cuts the *APOL1* gene sequence 74 bp downstream to the HindIII site (Figure S2A).

*APOL1* gene has two closely related paralogs, *APOL2* and *APOL3*, that could in principle interfere with library preparation (Figure S2B). To clear *APOL2* sequences, we took advantage of the fact that its sequence contains a recognition site for the NdeI restriction enzyme 36 bp upstream to the PvuII site and used this enzyme together with HindIII and PvuII. Although *APOL3* gene fragments could not be removed in a similar way since they do not contain any similar restriction sites, because they also lack the PvuII site, they could not undergo primary barcode addition and subsequent amplification alongside the APOL1 gene fragments. Additionally, the annealing site for the secondary barcode primer (APC) was selected to minimize hybridization to *APOL2* and *APOL3* sequences (Figure S2A).

Following enzymatic digestion and addition of the primary barcode sequences to the target DNA strand, a series of linear amplification reactions were used to create multiple copies of each barcoded molecule, with each copy being tagged by a unique secondary barcode (Figure S1A). This allowed us to distinguish at the sequence analysis step between de novo mutations and errors that were introduced by the DNA polymerase or the NGS machinery. Unlike such errors that tend to appear mostly in a single copy of the original sequence, true de novo mutations can be identified by their high frequency in groups of reads that share the same primary barcodes and their association with multiple secondary barcodes, which reflects their independent appearance in multiple copies made directly from the original target molecule.

##### 1.1.2 *APOL1* MEMDS consists of two reactions that run in parallel enabling accurate calculation of de novo mutation rates

An essential requirement for the calculation of de novo mutation rates is determining the number of WT (non-mutated) ROI-carrying molecules that are digested by HindIII during the mutation enrichment step. To accomplish this, we ran the MEMDS experiment using two parallel protocols, as depicted in Figure 1A (for a detailed description of this experimental design, see Melamed et al., 2022 [*1*]). Briefly, the two protocols are identical in all steps except the HindIII treatment, which is applied only to one of the two samples. We will refer to the samples as the “HindIII-treated” and “HindIII-untreated”, respectively. Artificial *APOL1* fragments, each carrying a known stretch of mutations instead of the HindIII site and therefore being resistant to the HindIII digestion are added in known amounts to each of the two samples prior to restriction enzyme treatment. At the sequence analysis step, the ratio of artificial HindIII-resistant to HindIII-susceptible (genomic, WT) sequences following each treatment is calculated. Dividing the ratio obtained for the “HindIII-treated” sample by the ratio obtained for the “HindIII-untreated” sample determines the HindIII-enrichment factor and consequently the total number of scanned genomes.

#### 1.2 Analysis of *APOL1* mutation variants from sperm samples of African and European donors

##### 1.2.1 Generating consensus sequences

Using the MEMDS procedure, we generated and sequenced DNA libraries from human sperm DNA of 7 African (AFR) and 8 European (EUR) donors (see Table S1 for donors' metadata). Prior to library preparations, all donors were verified for being homozygous to G0 (that is, having no G1 or G2 allelic variants) and for lacking any other DNA mutations that may interfere with library preparation, such as DNA mutations at primer annealing sites. At the sequence analysis step, high quality reads originating from direct extension of the target DNA strand were aligned against the *APOL1* reference sequence for the identification of mutations and grouped into "families" based on their primary barcode sequences (see Figure S1A, step 2, and Figure S3A). For each family, the consensus sequence that represents the sequence of the original target molecule was determined using three criteria (Figure 1A, step 8): the mutation frequency, which is the frequency of each mutation or a WT base within a family; the number of secondary barcodes, which is the number of unique secondary barcodes associated with each mutation or a WT base in the family, and family size, which is the total number of reads in the family. Both WT and mutated bases were filtered using the same criteria to eliminate bias in base calling (see also Figure S9 in Melamed et al., 2022 [*1*] for a detailed description of the algorithm). Finally, we separated the identified consensus mutations into two groups based on whether they are in the ROI site or in the sequences that flank the ROI (Figure S2A). Since mutations outside of the

ROI site are unlikely to be enriched by the HindIII treatment, we took the most stringent approach and considered their presence as being the result of library preparation or NGS errors. Consequently, the mutation rate inferred from the flanking sequences served as the basal false positive rate (FPR) of the MEMDS approach prior to its improvement by the HindIII-mutation enrichment step.

In accordance with previous studies [*1, 3*], we found that increasing the mutation frequency threshold and the number of secondary barcodes that were associated with each mutation or a WT base in a family increased the accuracy of mutation detection (Figure S4A). We found that mutation frequency $\geq 0.7$, number of secondary barcodes $\geq 3$ and family size $\geq 4$ provide an optimal balance between effective removal of false positives and loss of good data due to increased threshold stringency, as elaborated below.

Increasing the mutation frequency cutoff above 0.7 resulted in a notable loss of de novo mutation variants relative to WT sequences (Figure S4B), probably due to infiltration of chimeric WT reads to mutation variant families. The opposite scenario, in which sequences of mutation variants infiltrate into WT families is much less likely, due to the tiny amount of these variants in the samples. Increasing the secondary barcode threshold above 3 led to a strong reduction in the number of approved families while having only a minor effect on accuracy (Figure S4C). Under the conditions of mutation frequency cutoff of 0.7 and secondary barcode cutoff of 3, the minimal number of reads in a family had a minor effect on accuracy (Figure S4D). We therefore combined the above criteria with a minimal family size of at least 4 reads for further analysis.

The application of the combined cutoff criteria resulted in the approval of ~75% and

~55% of the families, on average, in the HindIII-treated and HindIII-untreated samples, respectively. The difference in the fraction of approved families between the two treatments was due to the smaller family sizes in the HindIII-untreated samples, which stemmed from a higher number of undigested families being captured by MEMDS. This resulted in more families that failed to meet the secondary barcode and family size threshold criteria (see Table S2 for a summary of the approved and filtered families).

##### 1.2.2 False positive rates in the flanking sequences

For both the HindIII-treated and HindIII-untreated samples, 96%-99% of the mutations in the flanking sequences that passed the cutoff criteria were either C→A or G→A in the sequenced strand, which corresponded to G→T and C→T in the target, labeled DNA strand (Figure S6). These two mutation types frequently arise both in vivo and in vitro from oxidized guanine that forms 8-oxoG and from deaminated cytosine or 5-methylcytosine that form uracil or thymidine, respectively. Due to their ubiquitous nature, these mutations are usually removed from the analysis during single strand–based consensus sequencing. Following previous literature, we removed them from consideration as de novo mutations in the ROI and from the FPR calculations [*1, 3, 4*].

After cleaning the G→T and C→T mutations, the FPR values per donor were calculated as described below. Averaged across all donors, the FPR obtained for non G→T, C→T mutations in the flanking sequences using the chosen cutoff criteria was $1.2 \times 10^{-6}$ ($\pm 0.5 \times 10^{-6}$) per base for the HindIII-treated samples and $2.5 \times 10^{-7}$ ($\pm 1.4 \times 10^{-7}$) per base for the HindIII-untreated samples (Figure S7). In accordance with the previous report of the FPRs obtained for the HBB and HBD genes using MEMDS [*1*], the FPR in the HindIII-treated samples is slightly higher than the one observed for the HindIII-untreated samples. The reason for this difference is likely the secondary influence of the large amount of processed DNA in the HindIII-treated samples compared to the HindIII-untreated samples (5% of the amount of genomic DNA used for the HindIII-treated sample is used for the HindIII-untreated sample) and also, possibly, the minor enrichment of mutations outside the HindIII-recognition site that may display weak inhibitory effects on HindIII. Yet, the high correlation between the frequencies of individual mutations in the flanking sequences of the HindIII-treated and HindIII-untreated samples suggests that the noise distribution for the two treatments is positively correlated ($R^2 = 0.66$, Fig. S8; Spearman's rank correlation: $\rho = 0.7$, $p = 2.14 \times 10^{-18}$). Additionally, we found that both the total FPR and the per-mutation-type FPR in the flanking sequences are highly similar between the African and European donors (Figures S7, S9), suggesting that any difference in mutation rate within the ROI site between the two populations cannot be assigned to differences in FPRs between the two groups.

##### 1.2.3 Enrichment factor of HindIII-resistant mutations and false positive rate in the *APOL1* ROI

Based on the changes in frequencies of the two artificial ROIs that were spiked into the HindIII-treated and HindIII-untreated samples, as described above, the calculated per sample HindIII enrichment score ranged from about 40 to 160, which represents approximately 97.5% to 99.5% digestion of WT ROI fragments by HindIII, respectively (Table S3). The average enrichment score for the African samples was 59.9 (±20.2), which represents removal by HindIII of 98.33% of the WT APOL1 sequences. The average enrichment score for the European samples was 100.2 (±44.9), which stands for removal of 99.02% of the WT APOL1 sequences.

We used the more conservative, higher, false positive rate obtained for the flanking sequences of the HindIII-treated samples as the baseline for the calculation of the false positive rate within the ROI in these samples. To obtain the FPR within the ROI for a specific sample, we divided the FPR obtained from the flanking sequences by the enrichment factor of this sample (see below and Figure S1B). Based on the per-individual enrichment factor, the average FPR calculated for the 6 bp of the ROI was ~$1.8 \times 10^{-8}$ with highly similar values between the two donor groups (Figure S7). Note that this value represents the upper bound of the expected noise, since we used the most conservative values of FPR in the flanking sequences to calculate the FPR within the ROI. The total number of scanned ROI fragments (i.e., fragments that were either sequenced or digested by HindIII) was calculated to be ~614 million: 291 million from the African and 323 from the European donors (Tables S1 and S3).

##### 1.2.4 Eliminating additional noise from the ROI of HindIII-treated samples

Of the non G→T non C→T mutations found in the ROI sequence of the HindIII-treated samples, three mutations, 1026C→A, 1027T→A and 1028T→A, were found to occur also in the ROI sequences of the HindIII-untreated samples. With the exception of these three mutations, no other mutation, including the 1024A→G G1 mutation, was found in the

ROI sequences of the HindIII-untreated samples across all donors. Though the absolute rates of these mutations were lower in the HindIII-untreated than in the HindIII-treated samples, adjusting them by the fold-difference in FPRs of the two treatments resulted in very similar mutation rates (Figure S10). Therefore, we considered these mutations to be errors introduced during library generation of both the HindIII-untreated and the HindIII-treated samples and excluded them from further analysis.

###### 1.2.5 Calculation of false positive rates and yields

The false positive rate of the flanking sequences per donor $i$, $\varepsilon'_i$, for either the HindIII-treated or untreated samples, is

$$\varepsilon'_i = \frac{M'_i}{F_i \times K'} \tag{1}$$

where $M'_i$ is the sum of all non G$\rightarrow$T non C$\rightarrow$T mutations in the flanking sequences of the target DNA strand across all families of donor $i$, $F_i$ is the number of approved sequence families of donor $i$, and $K'$ is the effective total number of bases across the flanking sequences in a DNA fragment (the number of different possible non G$\rightarrow$T non C$\rightarrow$T mutations across the flanking sequences in a fragment, divided by 3).

Calculating $\varepsilon'_i$ based on the HindIII-treated samples, the ROI FPR for each donor $i$, $\varepsilon_i$, is essentially equal to

$$\varepsilon_i = \frac{\varepsilon_i'}{E_i} \tag{2}$$

where $E_i$ is the HindIII enrichment factor for donor $i$. The per individual enrichment factor was calculated using the formula given in Figure 1A and its values are presented in Table S3.

The MEMDS yield per donor for flanking sequences, $Y_i'$, for either the HindIII-treated or untreated samples, is

$$Y_i' = \frac{F_i \times L'}{P_i'} \tag{3}$$

where $L'$ is the number of bases in the flanking sequences and $P_i'$ is the total number of bases that were sequenced in the flanking sequences of donor $i$.

The yield per donor for the ROI, $Y_i$, is essentially equal to

$$Y_i = \frac{F_i \times L \times E_i}{P_i} \tag{4}$$

where $L$ is the number of bases in the ROI and $P_i$ is the total number of bases that were sequenced in the ROI of donor $i$, for the HindIII-treated samples. Note that because the enrichment step enables detection of most WT ROIs without the need to sequence them first, the yield for the ROI can be higher than 1.

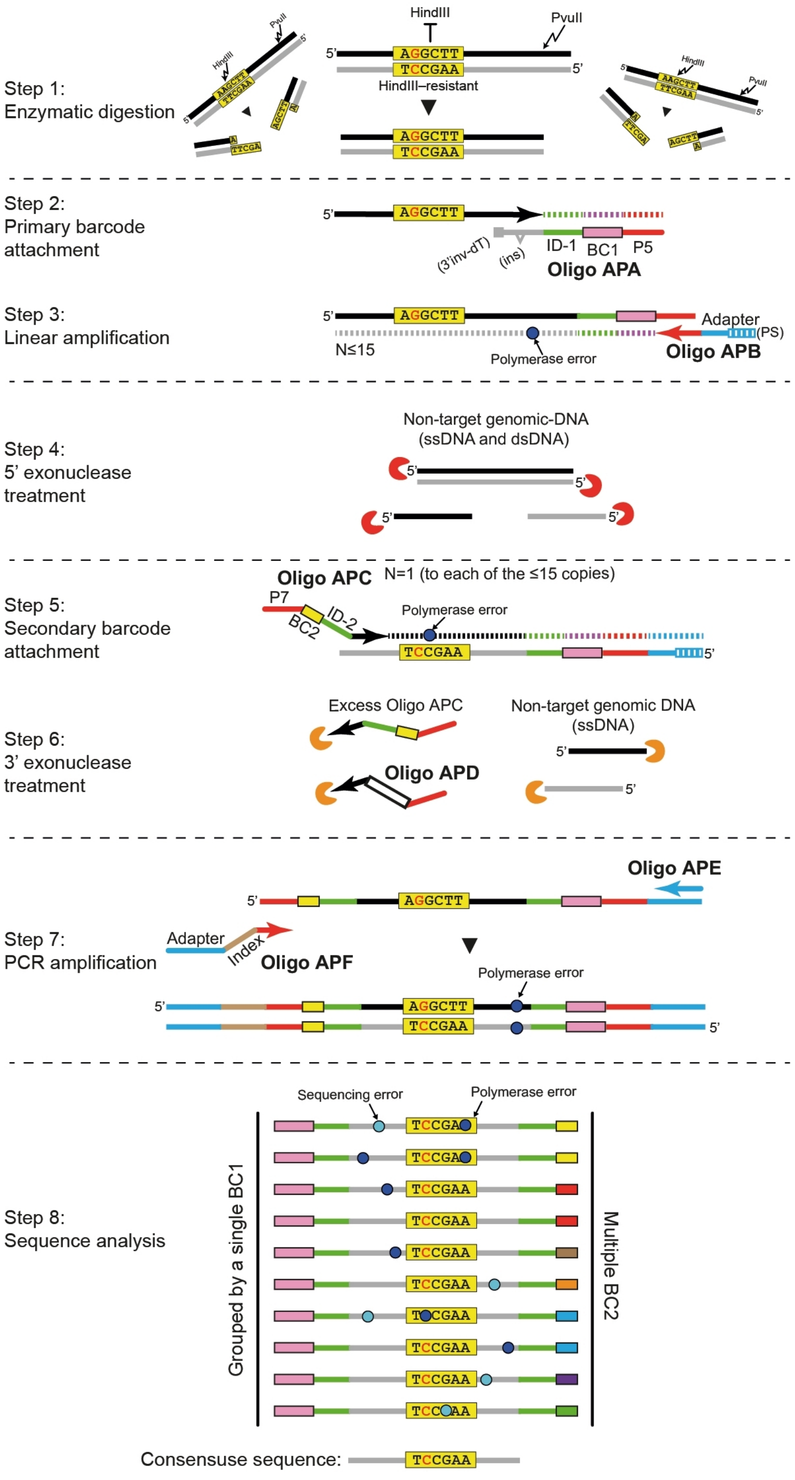

Step 1:
Enzymatic digestion
HindIII
PvuII
AGGCTT
TCCGAA
HindIII–resistant
Step 2:
Primary barcode attachment
(3'inv-dT)
(ins)
ID-1
BC1
P5
Oligo APA
Step 3:
Linear amplification
Adapter
(PS)
Oligo APB
N≤15
Polymerase error
Step 4:
5' exonuclease treatment
Non-target genomic-DNA
(ssDNA and dsDNA)
Oligo APC
N=1 (to each of the ≤15 copies)
Step 5:
Secondary barcode attachment
P7
BC2
ID-2
Polymerase error
Step 6:
3' exonuclease treatment
Excess Oligo APC
Non-target genomic DNA
(ssDNA)
Oligo APD
Oligo APE
Step 7:
PCR amplification
Adapter
Index
Oligo APF
Polymerase error
Step 8:
Sequence analysis
Sequencing error
Polymerase error
Grouped by a single BC1
Multiple BC2
Consensuse sequence:

**Figure S1: Applying MEMDS to study the de novo rates of the *APOL1* G1 1024A→G and nearby mutations – schematic illustration.** Dashed lines between steps represent DNA purification. **Step 1)** Sperm genomic DNA is digested by HindIII and PvuII. HindIII cuts wild type (WT) *APOL1* sequences at the AAGCTT restriction site that overlaps the region of interest (ROI) while leaving intact APOL1 sequences that carry the G1 1024A→G mutation at the ROI (AGGCTT) or other ROI HindIII-resistant mutations. This step results in enrichment of ROI mutations relative to WT sequences. **Step 2)** Sample identifier (ID-1), primary barcode (BC1) and Illumina P5 sequences are added directly to the original *APOL1* sense-DNA strand at the PvuII cleavage site by a fill-in reaction using Oligo APA (Table S4), which anneals to the sense strand and serves as a template. The extension of Oligo APA itself is blocked by a 3'-inverted dT (3'inv-dT) modification. Although we find this modification to be extremely efficient in blocking DNA polymerase Q5, a single-base thymine insertion (ins) in Oligo APA enables the identification of reads resulting from erroneous Oligo APA extensions and their removal at the sequence analysis step. **Step 3)** A linear amplification reaction is carried out by Oligo APB for 15 cycles. It generates multiple single-strand copies of barcoded *APOL1* sequences that can be grouped at the sequence analysis step by their shared primary barcode sequence. Since these copies are generated independently from each other, polymerase errors (dark blue circle) are unlikely to occur repeatedly in multiple copies originating from the same barcoded molecule as opposed to true de novo mutations. Oligo APB anneals to the Illumina P5 sequence and adds to each amplified copy an Illumina adapter sequence and five phosphorothioate bonds at its 5' terminus. **Step 4)** A mix of three 5' exonucleases is added to degrade the large amount of unrelated genomic DNA. The linearly amplified copies are protected from 5' exonucleases due to the phosphorothioate bonds at their 5' termini. **Step 5)** A single-cycle extension reaction is performed using Oligo APC, which adds sample identifier (ID-2), secondary barcode (BC2) and Illumina P7 sequences to the copied strands, with the secondary barcode being unique for each of the linearly amplified copies originating from the same target molecule (and hence carrying the same primary barcode sequence). **Step 6)** A 3' exonuclease is added to degrade excessive, unbound Oligo APC to reduce secondary-barcode relabeling events during the subsequent PCR reaction. To estimate the degree of erroneous relabeling due to inefficient Oligo APC removal, a known amount of relabeling control oligo (Oligo APD) is added to the 3' exonuclease reaction. Oligo APD is identical to Oligo APC with the exception that ID2 and BC2 are replaced by a known sequence. Identification of Oligo APD sequence signature at the sequence analysis step points to imperfect primer removal and serves as a proxy for the level of relabeling by Oligo APC. **Step 7)** A PCR reaction generates the final amplicon sequence. Polymerase errors are shared primary barcode sequences to determine the consensus sequence of the original molecule. Mutations due to DNA polymerase (dark blue circles) and sequencing (light blue circles) errors are filtered out based on their low frequencies in families and their lack of association with multiple secondary barcodes, whereas true de novo mutations such as the *APOL1* G1 1024A→G mutation are identified due to their high frequencies in families and association with multiple secondary barcodes. Note that since the sequencing output refers to the antisense strand, mutations that were present at the barcoded, sense strand are the reciprocals of the sequenced mutations.

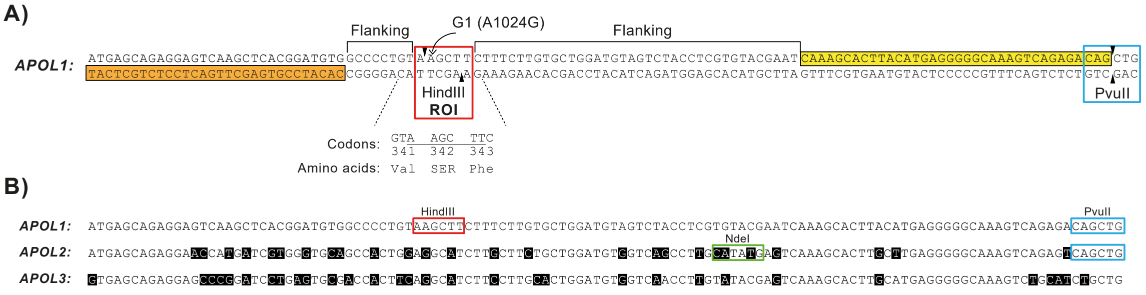


**Figure S2: APOL1 sequence features. A)** Shown is an *APOL1* DNA segment of 121 bp containing all of the gene features required for the MEMDS library generation. The upper sequence, which is in the sense orientation, serves as the target DNA strand, which is barcoded and subsequently amplified by the MEMDS protocol. The HindIII (RE-1)-recognition sequence appears in the red box and its cleavage sites are marked by small black triangles. The region of interest (ROI) overlaps with the HindIII site. A curved arrow points at position 1024, where the G1 1024A→G mutation occurs. The codon numbers and the amino acids encoded by the ROI sequence are shown underneath. The PvuII (RE-2)-recognition sequence appears in a blue box and its cleavage sites are marked by small black triangles. The sequence highlighted in yellow anneals to Oligo APA and receives the primary barcode via a single fill-in reaction (Figure S1A, step 2). The sequence highlighted in orange anneals to Oligo APC and receives the secondary barcode via a single extension reaction (Figure S1A, step 5). The sequence between Oligo APA and Oligo APC remains untouched by any primer and therefore is suitable for mutation detection analysis. However, only mutations at the ROI can be enriched, while mutations in the flanking sequences are unlikely to affect HindIII digestion. **B)** Sequence alignment of the two *APOL1* paralogs, *APOL2* and *APOL3*, to the *APOL1* library segment. Differences from the *APOL1* sequence are shown in black. Recognition sites for HindIII and PvuII appear in red and blue boxes, respectively. The recognition site for NdeI, which was used to eliminate *APOL2* sequences prior to library generation, appears in a green box. Additionally, the site for Oligo APC annealing (highlighted in orange in panel A) was chosen to minimize cross-reactivity with *APOL2* and *APOL3*. Yet, *APOL3* sequences were found in the *APOL1* sorted data in very small fractions (Table S2) and were removed prior to analysis. Note, however, that all of the ROI mutations that were detected in *APOL1* appeared as single base substitutions (or single base deletion) and were not associated with any other mutation in the ROI or the ROI flanking sequences, which rules out any possibility for their generation through *APOL1*-*APOL3* or *APOL1*-*APOL2* chimerism.

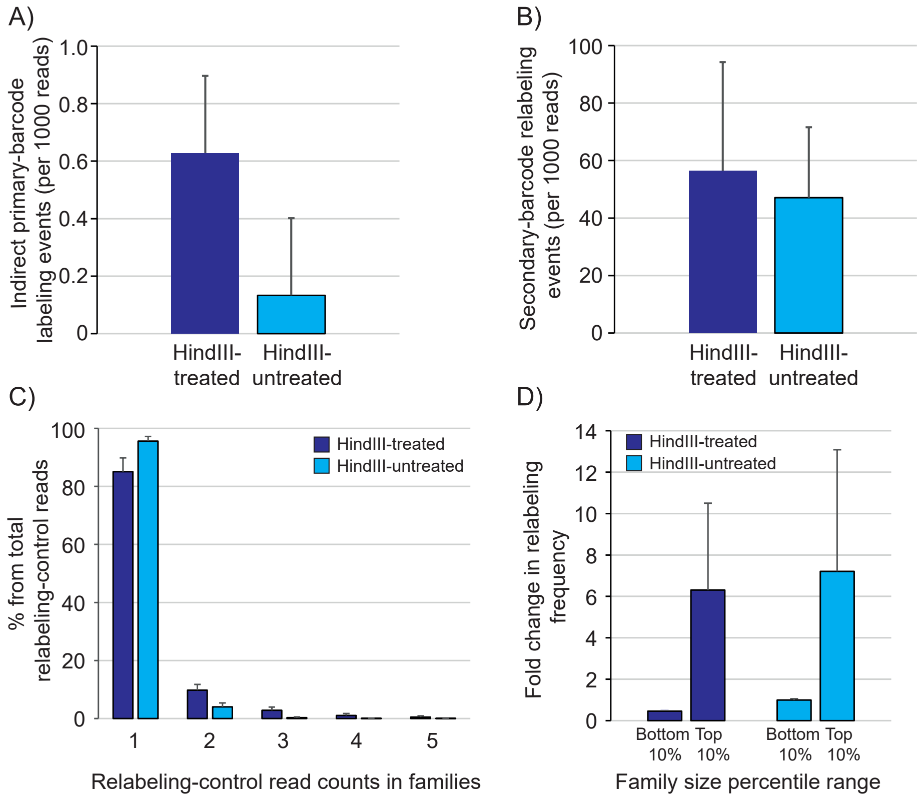


**Figure S3: Statistics of indirect barcode labeling and relabeling events at primary and secondary barcodes across the studied samples.** All values represent averages across all donors, calculated for each treatment (HindIII-treated and HindIII-untreated). **A)** Frequency of indirect labeling by the extension of the 3'-inverted dT blocked primary barcode oligo (Oligo APA), averaged across all samples, as measured by the fraction of reads carrying the control thymine insertion (Figure S1, step 2). **B)** The frequency of secondary barcode primer (Oligo APC) relabeling, averaged across all samples, as measured by the relative frequency of reads carrying the sequence signature of the control–secondary barcode relabeling primer (Oligo APD). **C)** Percent distribution of the number of relabeling events in families, using only families that have relabeling events (i.e., that carry the Oligo APD sequence signature). **D)** Fold-change in Oligo APD sequence signature frequencies in the top and bottom 10% of family sizes compared to its average frequency across all family sizes. Shown are average values for families with at least two reads (Family size $\geq$ 2) across all samples. Overall, relabeling events occur more frequently in larger families, but only in one or two reads per family, and thus are unlikely to affect the consensus analysis under the cutoff criteria used.

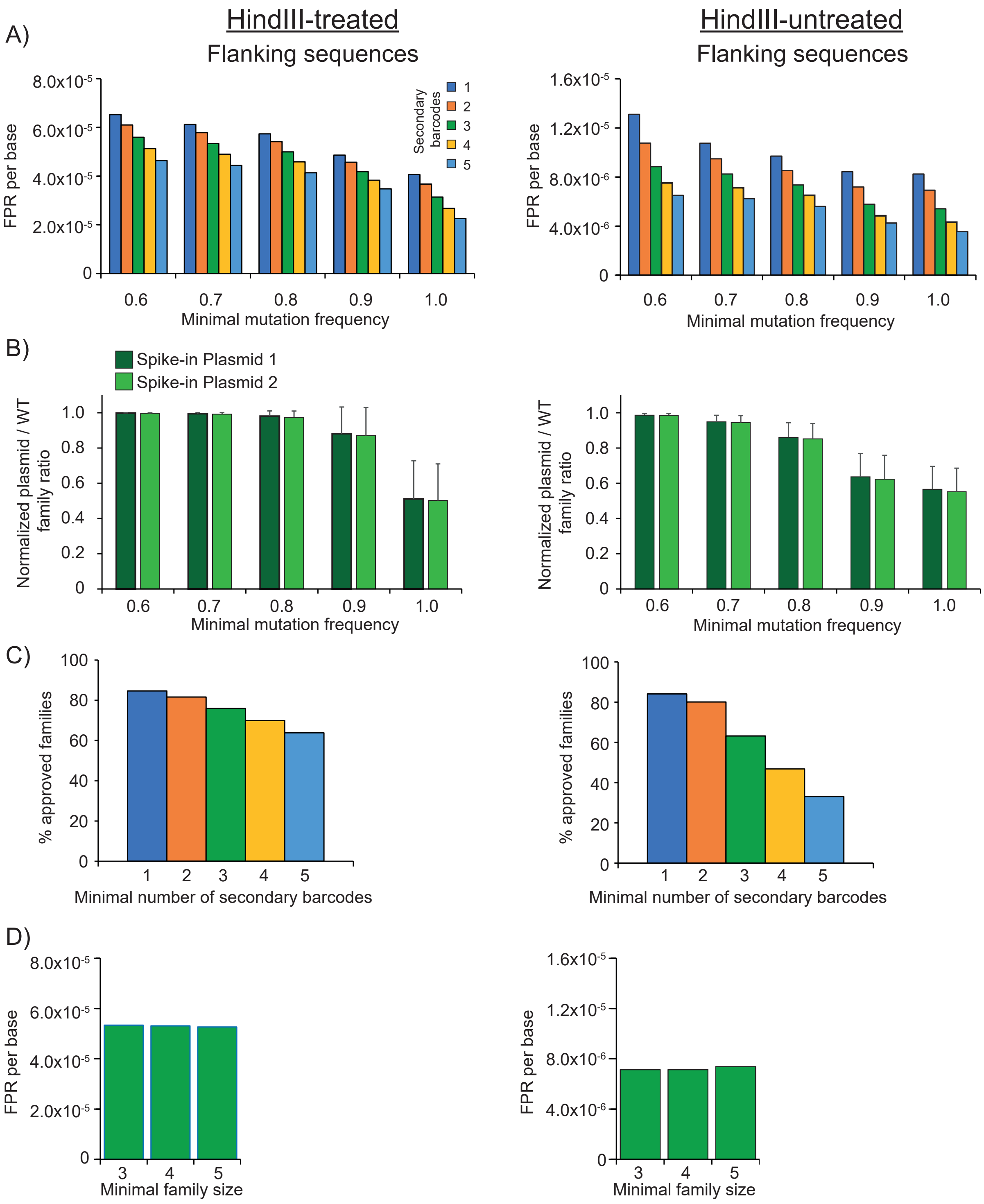
A)
HindIII-treated
Flanking sequences
FPR per base
8.0x10-5
6.0x10-5
4.0x10-5
2.0x10-5
0
Secondary barcodes
1
2
3
4
5
0.6
0.7
0.8
0.9
1.0
Minimal mutation frequency
HindIII-untreated
Flanking sequences
1.6x10-5
1.2x10-5
8.0x10-6
4.0x10-6
B)
Spike-in Plasmid 1
Spike-in Plasmid 2
Normalized plasmid / WT family ratio
1.0
0.8
0.6
0.4
0.2
C)
% approved families
100
80
60
40
20
Minimal number of secondary barcodes
D)
Minimal family size
3
4
5

**Figure S4: Mutation cutoff criteria.** Average values across all 15 donors for the HindIII-treated (left) and HindIII-untreated (right) samples informing the decision on the mutation cutoff criteria (mutation frequency, number of secondary barcodes and family size). **A)** Total FPRs for single-base substitutions, calculated across all single-base substitution types, including $C \rightarrow T$ and $G \rightarrow T$, in the flanking sequences, for families with at least two reads (family size $\geq 2$), using different combinations of minimal mutation frequency and secondary barcode cutoffs. **B)** The effect of mutation frequency cutoff (using family size $\geq 4$ and number of secondary barcodes $\geq 3$) on the spike-in–plasmid/WT family ratio. The plasmid/WT ratio was calculated for the two spike-in plasmids that were introduced to the HindIII-treated and HindIII-untreated samples for the calculation of HindIII-enrichment factor (Figure 1A). For each donor and treatment, the ratio was normalized to the ratio under minimal mutation frequency $\geq 0.5$. Note that for both the HindIII-treated and HindIII-untreated samples, increasing the mutation frequency cutoff above 0.7 results in a relatively stronger loss of plasmid variants (and presumably also natural mutation variants) compared to WT variants, probably due to infiltration of chimeric WT variants into the plasmid families (the opposite scenario, where plasmid sequences infiltrate and affect a large number of WT families is much less likely, due to the tiny amount of plasmid that is included in each sample). **C)** The effect of the minimal number of secondary barcodes that are associated with a mutated or a WT base on the number of approved primary barcode families (using family size $\geq 2$ and mutation frequency $\geq 0.7$). Specifically, a family is approved only if its size is larger than or equal to the family size cutoff, and if each position in the sequence contains a base (either a WT base or a mutated base) that meets both the mutation frequency and the secondary barcode cutoffs (otherwise the position is declared ambiguous and the family is rejected). The percent of approved families is determined relative to the combination of cutoff criteria that, by reducing ambiguity, allows the maximal number of families to pass filtration. Note that the HindIII-untreated sample shows a stronger secondary barcode cutoff effect on the percent of approved families, due to the smaller average family size in these samples. **D)** The effect of the family size cutoff on the FPR. Under mutation frequency $\geq 0.7$ and number of secondary barcodes $\geq 3$, the family size has a negligible effect on the FPR.

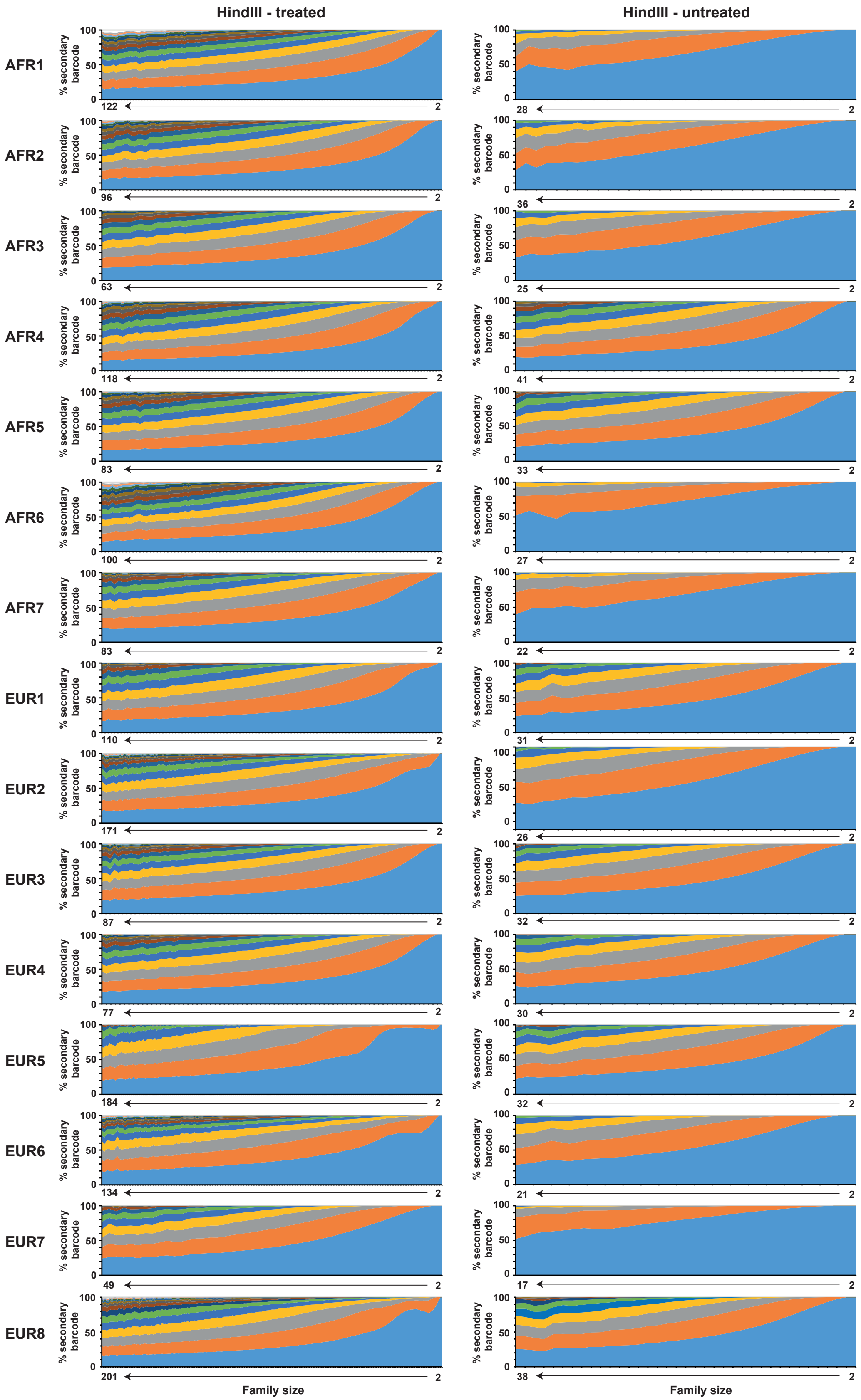
HindIII - treated
HindIII - untreated
AFR1
AFR2
AFR3
AFR4
AFR5
AFR6
AFR7
EUR1
EUR2
EUR3
EUR4
EUR5
EUR6
EUR7
EUR8
% secondary barcode
100
50
0
122
96
63
118
83
100
83
110
171
87
77
184
134
49
201
28
36
25
41
33
27
22
31
26
32
30
32
21
17
38
2
Family size

**Figure S5: The distribution of secondary barcodes in primary barcode families.** The figure shows the percentages of reads sharing the same secondary barcode in primary barcode families with respect to family size. Up to 15 unique secondary barcodes with at least two reads per barcode are shown. Their fractions were noted and averaged across all families of the same size. Only primary barcode families with at least two reads per family and family size groups with at least 10 families in them were analyzed. The noticeable differences in the maximal family sizes and in the secondary barcode distribution between the HindIII-treated and the HindIII-untreated samples were due to the large number of families obtained for the latter, which resulted in fewer reads per family. In accord with Figure S3 and the overall tendency of relabeling events to occur in only a few reads per family, in the vast majority of families the number of unique secondary barcode groups with more than one read in them does not exceed the expected number of linearly amplified copies (15).

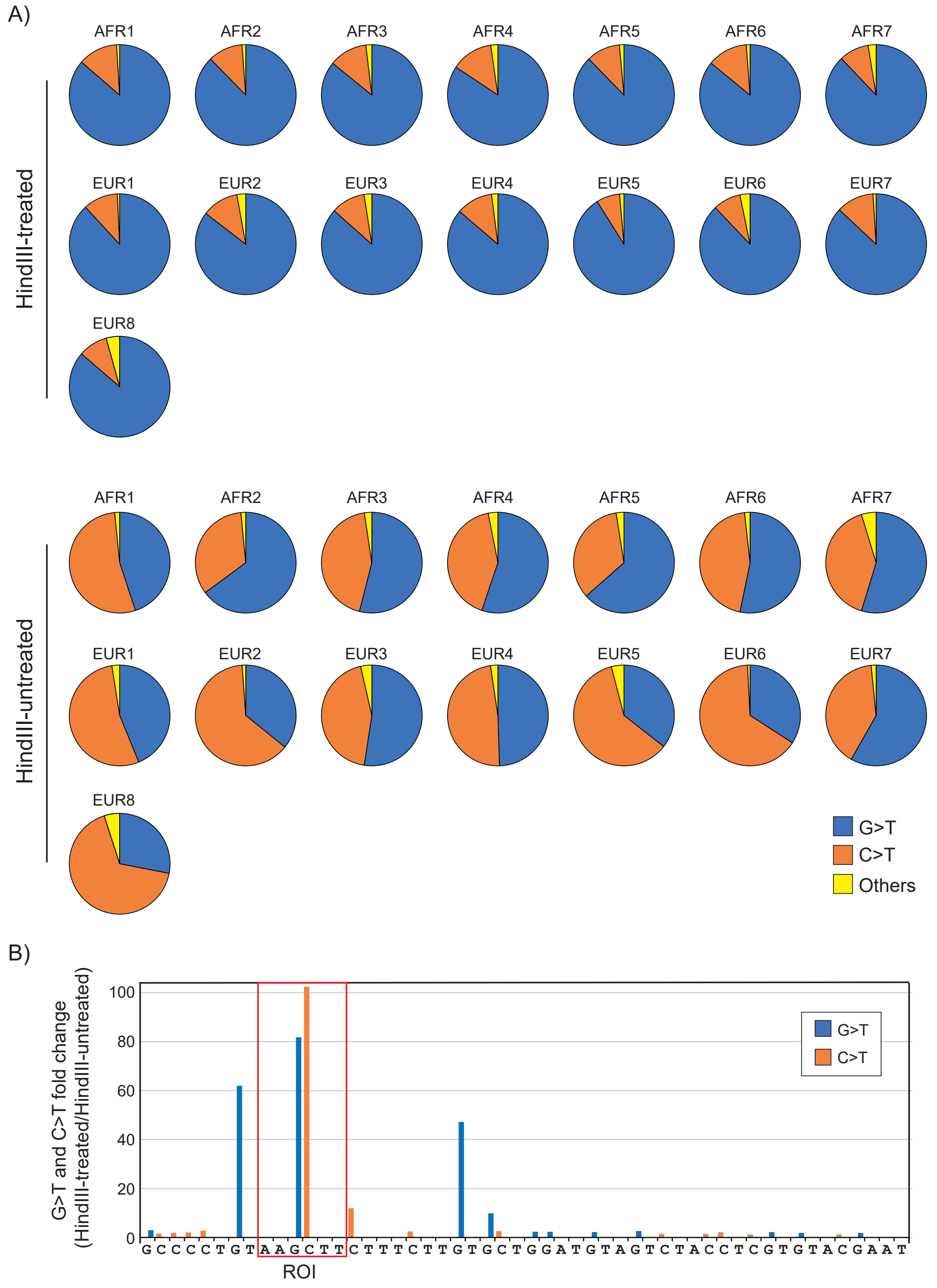
A)
HindIII-treated
AFR1
AFR2
AFR3
AFR4
AFR5
AFR6
AFR7
EUR1
EUR2
EUR3
EUR4
EUR5
EUR6
EUR7
EUR8
HindIII-untreated
AFR1
AFR2
AFR3
AFR4
AFR5
AFR6
AFR7
EUR1
EUR2
EUR3
EUR4
EUR5
EUR6
EUR7
EUR8
G>T
C>T
Others
B)
G>T and C>T fold change
(HindIII-treated/HindIII-untreated)
0
20
40
60
80
100
G>T
C>T
G C C C C T G T A A G C T T C T T T C T T G T G C T G G A T G T A G T C T A C C T C G T G T A C G A A T
ROI

**Figure S6: G→T and C→T are the most frequent point mutations. A)** Mutation type distributions in the flanking sequences of the *APOL1* target (sense) strand in the HindIII-treated and HindIII-untreated samples. **B)** HindIII-treated/HindIII-untreated ratios of G→T and C→T mutation frequencies per base. The source of the difference in the G→T and C→T distribution between the HindIII-treated and HindIII-untreated samples outside of the ROI (shown in panel A) can be attributed to a few specific positions in the analyzed reads. In particular, an approximately 60-fold increase in the G→T frequency in the HindIII-treated samples is observed two bases upstream to the ROI sequence (red box). Such enrichment upstream to a restriction site was also observed in the previous MEMDS-based study (Figure S13 in Melamed et al., 2022 [*1*]). Similarly, though of a smaller magnitude, a high C→T frequency in the HindIII-treated samples is observed one base downstream to the ROI sequence. In support to such enrichments, 8-oxoG modifications placed at restriction sites or near them have been shown to inhibit the activity of multiple restriction enzymes [*5–9*]. In addition, the formation of T:G or U:G mismatches due to deamination of cytosine or 5-meC have also been shown to interfere with enzymatic digestion [*10, 11*]. Finally, we observed an enrichment of G→T mutations eight and ten bases downstream to the ROI site, which may suggest a long-range destabilizing effect of guanine oxidation on the HindIII-restriction site.

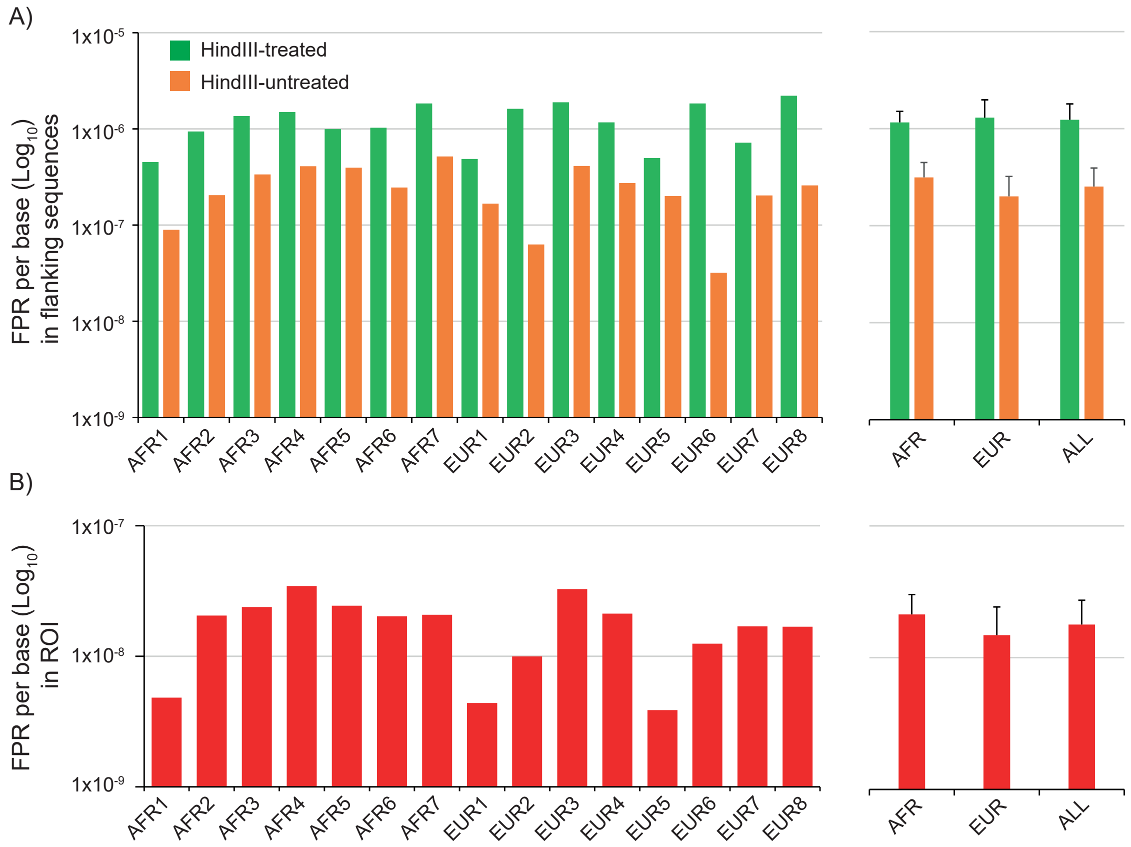


**Figure S7: False positive rates in the flanking sequences and ROI. A)** Non G→T, C→T FPRs in the flanking sequences of the HindIII-treated and HindIII-untreated samples. Per individual (left) and average (right) FPRs are shown. Higher FPRs for restriction enzyme-treated samples were observed previously [*1*] and were probably the result of the longer processing time required for these samples and/or the enrichment of mutations outside of the ROI that inhibit HindIII digestion, similar to the effects of some G→T and C→T mutations. For assessing the FPR in the ROI sequence of the HindIII-treated samples, we relied on the FPR in the flanking sequences of these samples, since it represents a more conservative estimate of noise in the ROI. **B)** Non G→T, C→T FPRs in the ROI sequence of the HindIII-treated samples, per individual (left) and averaged across African, European and all samples (right). The ROI FPR was calculated by dividing the FPR in the flanking sequences of each HindIII-treated sample by the HindIII-enrichment factor of that sample.

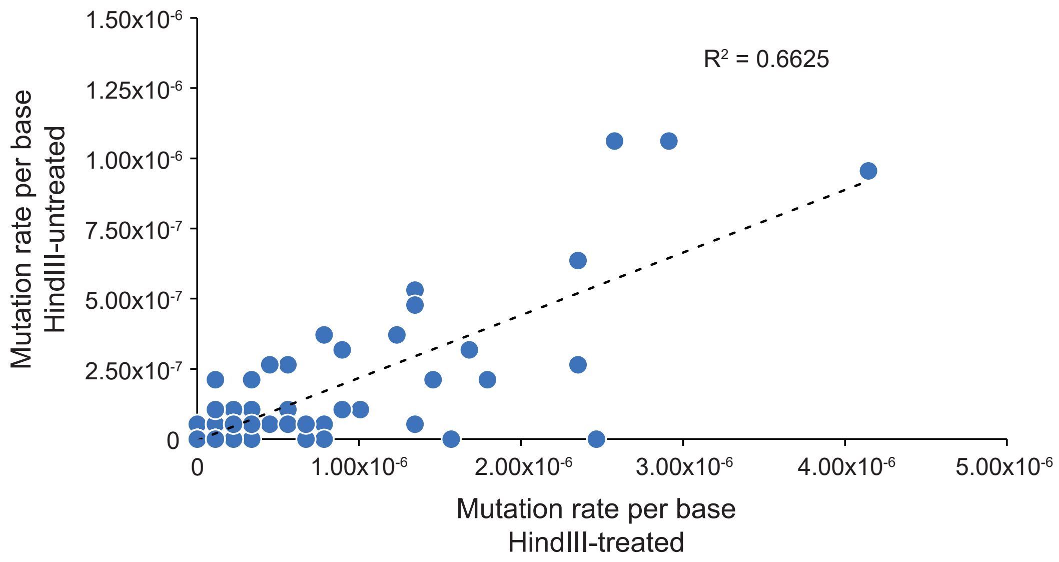


**Figure S8: Correlation between the rates of individual mutations in the flanking sequences of the HindIII-treated and HindIII-untreated samples.** Shown are rates of non G→T, C→T mutations based on summing up mutation instances across all donors separately for each treatment (HindIII-treated and HindIII-untreated). Each circle corresponds to a specific mutation at a specific position in the *APOL1* sequence.

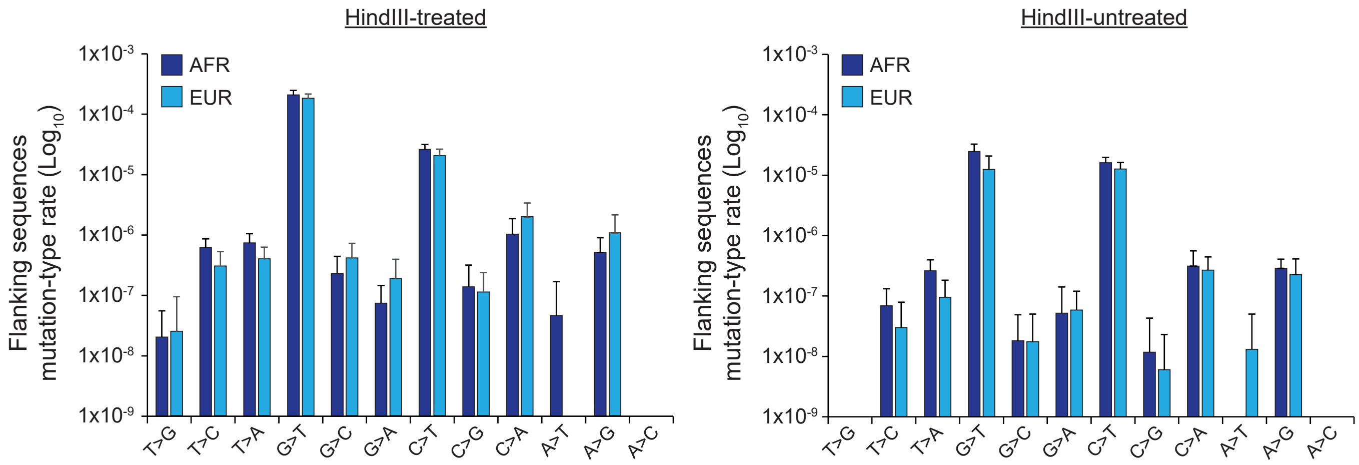


**Figure S9: Rates per mutation type in the flanking sequences, averaged across African (AFR) and across European (EUR) samples.** The figure shows mutation-type rates (including G$\rightarrow$T and C$\rightarrow$T) in the target (sense) DNA strand of the flanking sequences of the HindIII-treated (left) and HindIII-untreated (right) samples. The similarity between the populations in rates per mutation type in the flanking regions is consistent with the fact that the significant differences between the populations in mutation rates in the ROI are not due to differences in noise levels.

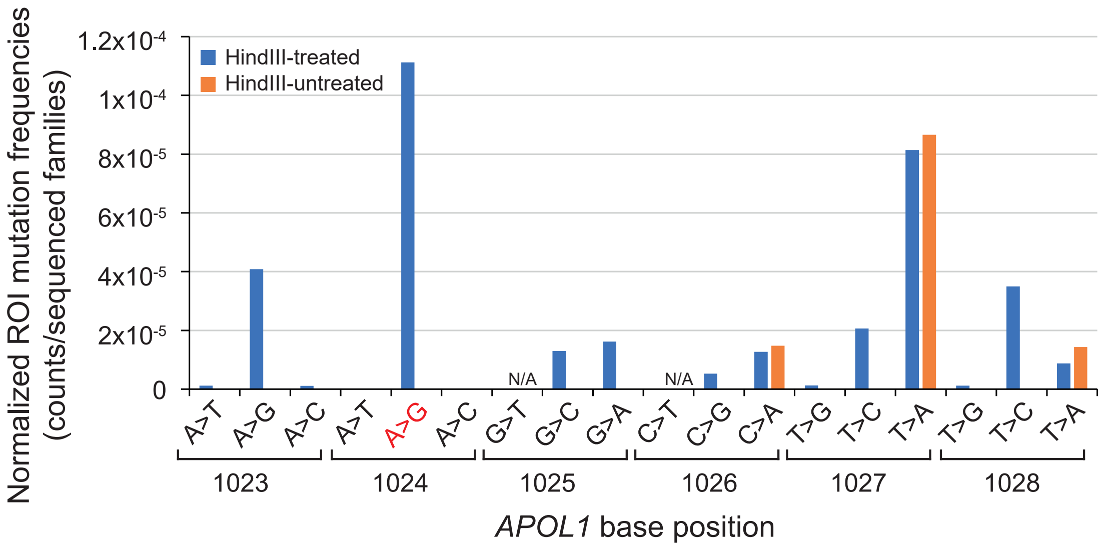


**Figure S10: Comparison of ROI mutation frequencies divided by the flanking sequences-based noise between the HindIII-treated and HindIII-untreated samples.** For each of the conditions (HindIII-treated and HindIII-untreated) we first calculated the mutation frequencies in the ROI as the total counts of ROI mutations across donors divided by the total number of families across donors (without considering enrichment). Because the flanking sequences–based FPR was higher in the HindIII-treated than in the HindIII-untreated samples, we then divided the ROI mutation frequencies of each condition by the flanking sequences–based average FPR of that condition. If any mutations observed in the ROI represent experimental artifacts (noise) rather than true de novo mutations, their noise-normalized rates are likely to be similar between the HindIII-treated and HindIII-untreated conditions. Indeed, we observed similar rates for the 1026C→A, 1027T→A and 1028T→A mutations for the HindIII-treated and untreated conditions and thus disqualified them from further analysis. Further supporting their exclusion, note that these three mutations are the only mutations that were found in the HindIII-untreated samples, while all the other mutations, including the *APOL1* G1 1024A→G (red) were observed only in the HindIII-treated samples. This fact supports the latter mutations' enrichment by the HindIII treatment and thus that the latter and not the former are true de novo mutations. G→T and C→T frequencies (reciprocals of C→A and G→A in the sequenced, non-labeled strand) are not included for the reasons discussed in the text.

**Table S1: Sperm samples' metadata**

| Donor | Donation year | Parent-1/ Parent-2 | Age group | Ethnic group | Haplotype | Sperm count (millions/mL) | % motility |
|---|---|---|---|---|---|---|---|
| **AFR1** | 2019 | Ghanaian/ Ghanaian | 23-28 | Akuapem | E150, I228, K255 | 58.9 | 45 |
| **AFR2** | 2018 | Ghanaian/ Ghanaian | 29-34 | Dagomba | E150, I228, K255 | 51.7 | 48 |
| **AFR3** | 2019 | Ghanaian/ Ghanaian | 29-34 | Ga | K150, I228, K255 | 75.3 | 62 |
| **AFR4** | 2019 | Ghanaian/ Ghanaian | 35-39 | Ewe | K/E150, I228, K255 | 47.1 | 42 |
| **AFR5** | 2018 | Ghanaian/ Ghanaian | 23-28 | Basari | K150, I228, K255 | 53.2 | 46 |
| **AFR6** | 2019 | Ghanaian/ Ghanaian | 35-39 | Effutu | E150, I228, K255 | 50.3 | 45 |
| **AFR7** | 2021 | Ghanaian/ Ghanaian | 18-22 | Ewe | K/E150, I228, K255 | 44.1 | 54 |
| **EUR1** | 2009/2011 | German/German | 18-22 | Caucasian | K150, I228, K255 | 79.1 | 67 |
| **EUR2** | 2001/2002/ 2003 | French/German | 18-22 | Caucasian | K/E150, I/M228, R/K255 | 87.3 | 71.6 |
| **EUR3** | 2012 | English/Irish-Italian | 18-22 | Caucasian | K150, I228, K255 | 90.5 | 55 |
| **EUR4** | 2011/2012 | Norwegian-Swedish/English-German | 18-22 | Caucasian | K150, I228, K255 | 90.1 | 55 |
| **EUR5** | 2005/6 | German/French-Italian | 23-28 | Caucasian | K/E150, I/M228, R/K255 | 57.2 | 55 |
| **EUR6** | 1994/1995 | English/Irish | 18-22 | Caucasian | K150, I228, K255 | 78.8 | 66 |
| **EUR7** | 2011/2012 | Polish/German-Irish-Scottish | 23-28 | Caucasian | K150, I228, K255 | 93.6 | 66 |
| **EUR8** | 2009 | English/English-French | 29-34 | Caucasian | K/E150, M228, K255 | 82.2 | 55 |

**Table S2: Counts of rejected and approved *APOL1* read families following filtration by the combined cutoff criteria**

| Donor | Treatment | Total Families [1] | Rejected: low size [2] | Rejected WT: low SB count [3] | Rejected: *APOL3* match [4] | Rejected: ambiguous (N) bases [5] | Approved Families [6] |
|---|---|---|---|---|---|---|---|
| **AFR1** | HindIII-treated | 763,683 | 146,728 | 6,743 | 1,262 | 2,691 | 606,259 |
| | HindIII-untreated | 2,811,294 | 897,231 | 10,936 | 5 | 39,148 | 1,863,974 |
| **AFR2** | HindIII-treated | 1,082,134 | 174,852 | 22,935 | 58 | 6,897 | 877,392 |
| | HindIII-untreated | 2,201,856 | 552,479 | 6,904 | 2 | 21,943 | 1,620,528 |
| **AFR3** | HindIII-treated | 1,155,547 | 216,610 | 30,125 | 144 | 8,768 | 899,900 |
| | HindIII-untreated | 2,783,422 | 1,268,362 | 39,124 | 0 | 27,414 | 1,448,552 |
| **AFR4** | HindIII-treated | 1,022,925 | 185,723 | 43,279 | 69 | 10,502 | 783,352 |
| | HindIII-untreated | 2,109,260 | 442,874 | 80,201 | 1 | 27,948 | 1,558,236 |
| **AFR5** | HindIII-treated | 1,204,030 | 155,070 | 76,139 | 29 | 11,528 | 961,264 |
| | HindIII-untreated | 1,979,486 | 686,009 | 150,746 | 0 | 32,901 | 1,109,830 |
| **AFR6** | HindIII-treated | 796,947 | 161,457 | 1,483 | 93 | 1,051 | 632,863 |
| | HindIII-untreated | 1,587,741 | 538,177 | 4,027 | 1 | 6,987 | 1,038,549 |
| **AFR7** | HindIII-treated | 726,460 | 144,258 | 45,265 | 75 | 8,593 | 528,269 |
| | HindIII-untreated | 2,483,389 | 1,391,088 | 22,857 | 1 | 13,990 | 1,055,453 |
| **EUR1** | HindIII-treated | 594,301 | 104,329 | 26,676 | 24 | 10,264 | 453,008 |
| | HindIII-untreated | 2,275,809 | 831,977 | 78,313 | 0 | 44,489 | 1,321,030 |
| **EUR2** | HindIII-treated | 490,053 | 121,649 | 5,711 | 113 | 2,540 | 360,040 |
| | HindIII-untreated | 2,938,930 | 1,146,359 | 31,693 | 0 | 27,754 | 1,733,124 |

**Table S2: Continued**

| Donor | Treatment | Total Families [1] | Rejected: low size [2] | Rejected WT: low SB count [3] | Rejected: *APOL3* match [4] | Rejected: ambiguous (N) bases [5] | Approved Families [6] |
|---|---|---|---|---|---|---|---|
| **EUR3** | HindIII-treated | 976,112 | 167,001 | 26,139 | 37 | 6,851 | 776,084 |
| | HindIII-untreated | 2,522,083 | 776,954 | 65,533 | 1 | 26,548 | 1,653,047 |
| **EUR4** | HindIII-treated | 911,442 | 154,379 | 59,989 | 190 | 12,406 | 684,478 |
| | HindIII-untreated | 2,398,036 | 894,228 | 78,493 | 1 | 26,855 | 1,398,459 |
| **EUR5** | HindIII-treated | 411,131 | 95,540 | 103,119 | 3 | 48,061 | 164,408 |
| | HindIII-untreated | 2,024,185 | 890,924 | 269,466 | 0 | 49,809 | 813,986 |
| **EUR6** | HindIII-treated | 406,709 | 73,902 | 5,843 | 21 | 1,588 | 325,355 |
| | HindIII-untreated | 2,349,780 | 1,402,985 | 80,187 | 0 | 21,551 | 845,057 |
| **EUR7** | HindIII-treated | 1,069,149 | 208,453 | 7,820 | 181 | 2,491 | 850,204 |
| | HindIII-untreated | 1,974,891 | 1,416,130 | 10,303 | 0 | 7,670 | 540,788 |
| **EUR8** | HindIII-treated | 449,345 | 129,239 | 8,819 | 204 | 2,986 | 308,097 |
| | HindIII-untreated | 2,540,152 | 690,807 | 52,661 | 1 | 16,585 | 1,780,098 |

1. Total number of families subjected to the combined cutoff criteria.
2. Families failing to meet the family size cutoff of 4.
3. Families consisting of non-mutated (complete wild-type) reads that meet the family size cutoff of 4 but fail to meet the secondary barcode count cutoff of 3.
4. Families with a consensus sequence that matches the *APOL3* gene fragment that was amplified by the MEMDS protocol due to its partial homology to the *APOL1* sequence (no record for *APOL2* families was found in the *APOL1* sorted library).
5. Mutation-containing families that fail to meet the mutation frequency cutoff of 0.7 or the secondary barcode count cutoff of 3 for either a mutation or a wild-type base in at least one of the positions in the sequence.
6. Families that passed the combined cutoff criteria of family size cutoff of 4, mutation frequency cutoff of 0.7 and secondary barcode count cutoff of 2. This number includes *APOL1* wild-type families, *APOL1* mutation families and spike-in plasmid families. Note that the numbers of all rejected and approved families sum up to the total number of families.

**Table S3: Values for the calculation of HindIII-enrichment factors and numbers of scanned target DNA sequences**

| Donor | Treatment | Donor DNA volume taken | Plasmid mix DNA volume taken | WT families passing cutoff [1] | Plasmid families passing cutoff [2] | Enrichment factor [3] | Scanned target sequences [4] |
|---|---|---|---|---|---|---|---|
| **AFR1** | HindIII-treated | 720 µl ($V_{Se}$) | 20 µl ($V_{Re}$) | 577,239 ($S_e^f$) | 14,753; 14,213 ($R_e^f$) | 93.49 | 53,968,355 |
| | HindIII-untreated | 36 µl ($V_{Sc}$) | 120 µl ($V_{Rc}$) | 1,750,829 ($S_c^f$) | 59,749; 53,016 ($R_c^f$) | 1 | 1,750,829 |
| **AFR2** | HindIII-treated | 360 µl ($V_{Se}$) | 20 µl ($V_{Re}$) | 855,765 ($S_e^f$) | 9,776; 8,529 ($R_e^f$) | 45.98 | 39,346,491 |
| | HindIII-untreated | 18 µl ($V_{Sc}$) | 120 µl ($V_{Rc}$) | 1,534,094 ($S_c^f$) | 45,694; 39,950 ($R_c^f$) | 1 | 1,534,094 |
| **AFR3** | HindIII-treated | 600 µl ($V_{Se}$) | 20 µl ($V_{Re}$) | 881,287 ($S_e^f$) | 8,414; 7,190 ($R_e^f$) | 57.00 | 50,235,510 |
| | HindIII-untreated | 30 µl ($V_{Sc}$) | 120 µl ($V_{Rc}$) | 1,395,717 ($S_c^f$) | 28,741; 23,283 ($R_c^f$) | 1 | 1,395,717 |
| **AFR4** | HindIII-treated | 770 µl ($V_{Se}$) | 20 µl ($V_{Re}$) | 764,485 ($S_e^f$) | 8,924; 7,550 ($R_e^f$) | 43.33 | 33,127,436 |
| | HindIII-untreated | 38.5 µl ($V_{Sc}$) | 120 µl ($V_{Rc}$) | 1,469,761 ($S_c^f$) | 47,497; 40,211 ($R_c^f$) | 1 | 1,469,761 |
| **AFR5** | HindIII-treated | 360 µl ($V_{Se}$) | 20 µl ($V_{Re}$) | 943,075 ($S_e^f$) | 8,274; 6,820 ($R_e^f$) | 40.75 | 38,433,605 |
| | HindIII-untreated | 18 µl ($V_{Sc}$) | 120 µl ($V_{Rc}$) | 1,059,254 ($S_c^f$) | 27,543; 22,377 ($R_c^f$) | 1 | 1,059,254 |
| **AFR6** | HindIII-treated | 350 µl ($V_{Se}$) | 20 µl ($V_{Re}$) | 607,891 ($S_e^f$) | 12,212; 10,285 ($R_e^f$) | 50.72 | 30,829,315 |
| | HindIII-untreated | 17.5 µl ($V_{Sc}$) | 120 µl ($V_{Rc}$) | 954,443 ($S_c^f$) | 46,140; 37,438 ($R_c^f$) | 1 | 954,443 |
| **AFR7** | HindIII-treated | 740 µl ($V_{Se}$) | 20 µl ($V_{Re}$) | 509,797 ($S_e^f$) | 9,072; 7,579 ($R_e^f$) | 88.25 | 44,991,174 |
| | HindIII-untreated | 37 µl ($V_{Sc}$) | 120 µl ($V_{Rc}$) | 1,010,732 ($S_c^f$) | 25,371; 19,517 ($R_c^f$) | 1 | 1,010,732 |
| **EUR1** | HindIII-treated | 380 µl ($V_{Se}$) | 20 µl ($V_{Re}$) | 428,297 ($S_e^f$) | 12,618; 10,645 ($R_e^f$) | 110.63 | 47,383,311 |
| | HindIII-untreated | 19 µl ($V_{Sc}$) | 120 µl ($V_{Rc}$) | 1,247,216 ($S_c^f$) | 40199; 33280 ($R_c^f$) | 1 | 1,247,216 |
| **EUR2** | HindIII-treated | 580 µl ($V_{Se}$) | 20 µl ($V_{Re}$) | 338,430 ($S_e^f$) | 10,967; 9,648 ($R_e^f$) | 162.07 | 54,849,563 |
| | HindIII-untreated | 29 µl ($V_{Sc}$) | 120 µl ($V_{Rc}$) | 1,657,992 ($S_c^f$) | 40,971; 33,807 ($R_c^f$) | 1 | 1,657,992 |
| **EUR3** | HindIII-treated | 450 µl ($V_{Se}$) | 20 µl ($V_{Re}$) | 758,365 ($S_e^f$) | 8,087; 6,763 ($R_e^f$) | 57.63 | 43,704,050 |
| | HindIII-untreated | 22.5 µl ($V_{Sc}$) | 120 µl ($V_{Rc}$) | 1,587,595 ($S_c^f$) | 35,332; 29,401 ($R_c^f$) | 1 | 1,587,595 |
| **EUR4** | HindIII-treated | 320 µl ($V_{Se}$) | 20 µl ($V_{Re}$) | 668,927 ($S_e^f$) | 7,380; 6,362 ($R_e^f$) | 55.05 | 36,824,698 |
| | HindIII-untreated | 16 µl ($V_{Sc}$) | 120 µl ($V_{Rc}$) | 1,337,939 ($S_c^f$) | 32,558; 27,356 ($R_c^f$) | 1 | 1,337,939 |
| **EUR5** | HindIII-treated | 640 µl ($V_{Se}$) | 20 µl ($V_{Re}$) | 157,185 ($S_e^f$) | 3,926; 3,000 ($R_e^f$) | 128.23 | 20,155,917 |
| | HindIII-untreated | 32µl ($V_{Sc}$) | 120 µl ($V_{Rc}$) | 781,602 ($S_c^f$) | 14,310; 17,919 ($R_c^f$) | 1 | 781,602 |
| **EUR6** | HindIII-treated | 790 µl ($V_{Se}$) | 20 µl ($V_{Re}$) | 311,813 ($S_e^f$) | 6,925; 5,689 ($R_e^f$) | 146.94 | 45,816,352 |
| | HindIII-untreated | 39.5 µl ($V_{Sc}$) | 120 µl ($V_{Rc}$) | 817,907 ($S_c^f$) | 15,105; 11,917 ($R_c^f$) | 1 | 817,907 |
| **EUR7** | HindIII-treated | 320 µl ($V_{Se}$) | 20 µl ($V_{Re}$) | 832,316 ($S_e^f$) | 8,237; 6,944 ($R_e^f$) | 42.39 | 35,284,884 |
| | HindIII-untreated | 16 µl ($V_{Sc}$) | 120 µl ($V_{Rc}$) | 513,995 ($S_c^f$) | 14,754; 11,783 ($R_c^f$) | 1 | 513,995 |
| **EUR8** | HindIII-treated | 510 µl ($V_{Se}$) | 20 µl ($V_{Re}$) | 294,591 ($S_e^f$) | 6,933; 5,927 ($R_e^f$) | 130.99 | 38,587,287 |
| | HindIII-untreated | 25.5 µl ($V_{Sc}$) | 120 µl ($V_{Rc}$) | 1,720,925 ($S_c^f$) | 32,466; 36,358 ($R_c^f$) | 1 | 1,720,925 |

1. Number of families with no mutations in the ROI that passed the combined cutoff criteria.
2. Number of families of spike-in plasmids that passed the combined cutoff criteria.
3. HindIII-enrichment factor, calculated from the volumes and family counts shown in the table using the formula shown in Figure 1A.
4. For the HindIII-treated samples, this number includes both the number of sequenced and the number of HindIII-digested WT target sequences, and is computed by multiplying the number of sequenced WT families by the enrichment factor. For the HindIII-untreated samples, this number is equal to the number of sequenced families.

**Table S4: Oligos used for library generation by the MEMDS protocol and for genotyping sperm donors**

| Oligos for MEMDS protocol | |
|---|---|
| **Name** | APA |
| **Used for** | Direct attachment of primary barcode |
| **Sequence** | P-*CTCTTTCCCTACACGACGCTCTTCCGATCT*(14N)XXXXCTGTCTCTGACTTTGCCtCCCTCATGTAAGTGCTTTG-InvdT |
| **Sequence features** | P – 5' phosphate, to improve degradation by 5' exonucleases. In italics – part of Illumina TruSeq Universal Adapter P5. (14N) – Primary Barcode, unique to each labeled molecule. XXXX – Donor identifier (ID)-1 composed of four bases unique to each treatment and each donor. Underlined sequence – complementary to APOL1 sense strand and start at the edge of PvuII digestion product. Lowercased "t" –a base insertion designed to identify events of erroneous extension and amplification promoted by unblocked (3' InvdT missing) Oligo APA, if any. InvdT – 3' inverted dT modification, designed to block extension by Q5 DNA polymerase. |
| **Name** | APB |
| **Used for** | Linear amplification of barcoded strand |
| **Sequence** | *ApsApsTpsGpsApsTACGGCGACCACCGAGATCTACA*CTCTTTCCCTACACGACGCTC |
| **Sequence features** | Underlined sequence – region complementary to the fill-in product (barcode-labeled strand) by Oligo APA. In italics – completes the 5' part of Illumina TruSeq Universal Adapter P5. ps – phosphorothioate; protects linearly amplified sequences from 5' exonuclease degradation. |
| **Name** | APC |
| **Used for** | Secondary-barcode labeling |
| **Sequence** | *GTGACTGGAGTTCAGACGTGTGCTCTTCCGATCT*(5N)XXXXATGAGCAGAGGAGTCAAGCTCACGGATGTG |
| **Sequence features** | In italics – Part of Illumina TruSeq Universal Adapter P7. (5N) – Secondary Barcode, labels each linearly amplified sequence that was generated by APB. XXXX – Donor identifier (ID)-2, composed of four bases unique to each donor. Underlined sequence – region complementary to APOL1 sequence. |
| **Name** | APD |
| **Used for** | Control for secondary-barcode relabeling |
| **Sequence** | *GTGACTGGAGTTCAGACGTGTGCTCTTCCGATCT*AGTGTAAAAATGAGCAGAGGAGTCAAGCTCACGGATGTG |
| **Sequence features** | Identical to Oligo APC, but with the sequence AGTGT replacing the Secondary barcode sequence (5N), and AAAA replacing ID-2 sequence XXXX (underlined) |
| **Name** | APE |
| **Used for** | Forward amplification primer for the first and the second PCR reactions |
| **Sequence** | AATGATACGGCGACCACCGAGATCTAC |
| **Sequence features** | Matches the 5' edge generated by APB |

**Table S4: Continued.**

| Oligos for MEMDS protocol | |
|---|---|
| **Name** | APF1 |
| **Used for** | Reverse amplification primer for the first PCR reaction |
| **Sequence** | GTGACTGGAGTTCAGACGTGTGCTC |
| **Sequence features** | Matches the 3’ edge generated by APC |
| **Name** | APF2 |
| **Used for** | Reverse amplification primer for the second PCR reaction |
| **Sequence** | CAAGCAGAAGACGGCATACGAGATXXXXXXGTGACTGGAGTTCAGACGTGTGC |
| **Sequence features** | Completes the Illumina TruSeq Universal Adapter P7; XXXXXX stands for the Illumina index sequence. |

| Oligos for APOL1 genotyping | | |
|---|---|---|
| **Name** | **Sequence** | **Used for** |
| **APOL1-F** | ACAAGCCCAAGCCCACGACC | Forward (F) and reverse (R) primers to validate G0 haplotype and compatibility of *APOL1* library sequence for MEMDS |
| **APOL1-R** | CCTGGCCCCTGCCAGGCATA | |
| **APOL1 E150-F** | CCTTTAACCTTTCCTTGTGCAG | Forward (F) and reverse (R) primers for validating *APOL1* gene background haplotype |
| **APOL1 E150-R** | CTGCTGGTAATCCCGGTCAA | |
| **APOL1 I228-F** | GTGGTGTCTGGCTCTCTCAG | |
| **APOL1 I228-R** | GCCTCGTGTGAGTTGGTAAGTA | |
| **APOL1 K255-F** | TGGAGTTGGGAATCACAGCC | |
| **APOL1 K255-R** | CTCCACCTGTTCACCGCTTT | |
